# Optimal cell length for exploration and exploitation in chemotactic planktonic bacteria

**DOI:** 10.1101/2023.10.05.560636

**Authors:** Òscar Guadayol, Rudi Schuech, Stuart Humphries

**Affiliations:** Mediterranean Institute for Advanced Studies, IMEDEA (UIB-CSIC), Miquel Marqués 21, 07190 Esporles, Balearic Islands, Spain; Department of Mathematics and Center for Computational Science, Tulane University, New Orleans, LA 70118, USA; School of Life Sciences, Joseph Banks Laboratories, University of Lincoln, Lincoln LN6 7DL, UK

**Author notes:** **Correspondence to:** Òscar Guadayol, Mediterranean Institute for Advanced Studies, IMEDEA (UIB-CSIC), Miquel Marqués 21, 07190 Esporles, Balearic Islands, Spain. **Author Contribution Statement OG**: Conceptualization, Methodology, Software, Formal analysis, Investigation, Data curation, Writing – Original Draft, Visualization. **RS**: Conceptualization, Methodology, Software, Writing-Review & Editing. **SH**: Conceptualization, Writing - Review & Editing, Supervision, Project Administration, Funding Acquisition.

## Abstract

Elongated morphologies are prevalent among motile bacterioplankton in aquatic systems. This is often attributed to enhanced chemotactic ability, but how long is best? We hypothesized the existence of an optimal cell length for efficient chemotaxis resulting from shape-imposed physical constraints acting on the trade-off between rapid exploration versus efficient exploitation of nutrient sources. To test this hypothesis, we first evaluated the chemotactic performance of elongated cephalexin-treated *Escherichia coli* towards α-methyl-aspartate in an agarose-based microfluidic device that creates linear, stable and quiescent chemical gradients. Our experiments showed that cells of intermediate lengths aggregated most tightly to the chemoattractant source. We then replicated these experimental results with Individual Based Model (IBM) simulations. A sensitivity analysis of the IBM allowed us to gain mechanistic insights into which parameters drive this trend and showed that the poor chemotactic performance of very short cells is caused by loss of directionality, whereas long cells are penalized by brief, slow runs. Finally, we evaluated the chemotactic performance of cells of different length with IBM simulations of a phycosphere – a hotspot of microbial interactions in the ocean. Results indicated that long cells swimming in a run-and-reverse pattern with extended runs and moderate speeds are most efficient at harvesting nutrients in this microenvironment. The combination of microfluidic experiments and IBMs proves thus to be a powerful tool for untangling the physical constraints that motile bacteria are facing in aquatic systems.

## Introduction

Shape is a functional trait that influences several important tasks in bacterial life, including motility, nutrient uptake, resistance to grazing, passive dispersion, and pathogenicity (Young 2006). Among the characters commonly used to parametrize microbial shape, cell length exhibits one of the highest degrees of variability. Not only is there considerable interspecific diversity in cell length but also intraspecific. Cell length may fluctuate throughout the cell cycle and depends on growth rate and nutritional environment (Vadia and Levin 2015; Willis and Huang 2017) and sometimes increases dramatically as stressed individuals stop dividing (Erill et al. 2007).

One of the most important behaviours that planktonic bacteria can undertake is chemotaxis, which allows a motile bacterium to detect chemical gradients (exploration) and to track these gradients and remain close to the nutrient source to maximize uptake (exploitation) (Stocker and Seymour 2012). Elongated cell morphologies are particularly prevalent among planktonic chemotactic bacteria (Dusenbery 1998) probably because they help microbial swimmers overcome the most insidious challenge they face: Brownian motion. Swimming bacteria live in a low Reynolds number regime and, due to their small size, cannot maintain a straight course as they are constantly deflected by colliding water molecules. To overcome this challenge, flagellated bacteria have evolved a characteristic swimming pattern, consisting of a sequence of runs and sudden reorientations. During a run, a cell swims in a relatively straight line, whereas in the reorientation phase the direction of motion changes in species-specific ways (Mitchell 2002; Mitchell and Kogure 2006; Grognot and Taute 2021).

Elongation, by increasing rotational drag, reduces the effects of Brownian motion, thus allowing longer and straighter runs and increasing signal-to-noise ratios and chemotactic performance (Dusenbery 2009). Furthermore, the increase in rotational drag due to elongation also restricts reorientation angles (Guadayol et al. 2017), which helps cells to track gradients more rapidly (Locsei 2007; Nicolau et al. 2009).

Elongation also poses some challenges to efficient chemotaxis. First, the restriction of reorientation angles leads to higher dispersal rates (Lovely and Dahlquist 1975) and so shortens the time spent close to a nutrient source. Second, elongation increases translational friction, slowing the cell and decreasing swimming efficiency (Dusenbery 1998; Schuech et al. 2019). And finally, it can slow down or weaken the cell’s response to environmental cues as, given the diffusive nature of intracellular transport of signals, the time it takes for chemotactic signals to travel from the membrane receptors to the flagellar motors increases quadratically with cell length (Segall et al. 1985). In bacteria with randomly distributed multiple flagella, this signal delay can also induce desynchronization of flagella (Maki et al. 2000), ultimately leading to alterations in the motility pattern (Guadayol et al. 2017).

In summary, current theories predict both potential advantages and disadvantages of elongation for chemotaxis (see *Supplementary material*, *Theoretical Background* and Table S1). We hypothesized that optimal cell lengths for efficient chemotaxis exist, imposed by a trade-off between the need to run straight, swim efficiently, and explore the environment quickly, and the need to reorient when desired, respond promptly to environmental changes, and maximize the time spent near nutrient sources.

We sought to experimentally test our hypothesis by using *Escherichia coli* cultures treated with cephalexin, an antibiotic that stops cells from dividing without otherwise physiologically impairing them (Rolinson 1980; Ishihara et al. 1983; Maki et al. 2000). To assess the influence of cell length on chemotactic performance, we subjected cultures at various stages of elongation to microfluidic chemotactic assays with different gradients of concentration of the chemoattractant α-methyl-aspartate. To gain a mechanistic understanding of the experimental results, we then built an individual based model (IBM) of our experimental system in which the chemotactic behaviour was simulated with a well-known model for the aspartate chemotactic pathway in *E. coli* (Tu et al. 2008). We used this IBM to further explore the consequences of elongation for chemotaxis in the phycosphere, the chemically distinct area surrounding phytoplanktonic cells where most interactions with bacteria, with global significance, take place (Seymour et al. 2017; Raina et al. 2022).

## Materials and methods

### Cultures

Experiments were performed with the chemotactic *E. coli* strain AW405. Cultures were grown overnight at 30°C in 25 mL of minimum growth media (Maki et al. 2000) from frozen glycerol stock and shaken at 200 rpm at an orbit of 19 mm in 100 mL Erlenmeyer flasks. The saturated culture was diluted 100-fold in 25 mL of fresh medium. Growth was monitored hourly using a cell density meter measuring at a wavelength of 600 nm. After ∼2 h, once cultures had achieved exponential growth (optical density ∼0.2), a sample was taken to conduct a chemotactic assay on the non-treated population, and then cephalexin was added to a final concentration of 60 µgL^-1^. Cephalexin is a β- lactam antibiotic that promotes the formation of increasingly elongated cells and filaments by interfering with the formation of the septal ring and stopping cell division (Eberhardt et al. 2003). Cephalexin is not otherwise known to alter growth rate, physiology, or flagellar motility (Rolinson 1980; Ishihara et al. 1983; Maki et al. 2000). Once cephalexin was added, chemotactic performance was evaluated every hour, as cells progressively elongated, up to five hours after the addition of cephalexin.

Before performing any chemotactic assay, cultures were washed into random motility buffer (RMB, Turner et al. 2000) by filtering and backwashing with RMB through a 0.2 µm sterile membrane three times (Bell et al. 2005). This procedure resulted in highly motile suspensions of bacteria with limited loss of cells (usually less than half) and was much quicker than the classical centrifugation technique.

### Microfluidic device

To evaluate the chemotactic ability of *E. coli,* we used an agarose-based microfluidic device with three channels (Fig. 1A, Diao et al. 2006; Cheng et al. 2007; Ahmed et al. 2010). This design allows the generation of quiescent, steady linear chemical gradients. We used a design with three 600 µm wide and 100 µm deep channels running parallel and separated by 200 µm wide walls and embedded in 3% agarose (corresponding to ‘design 2’ in Ahmed et al. 2010). Details on the fabrication and calibration of the devices are given in *Supplementary material*, *Microfluidic devices*.

**Figure 1:**
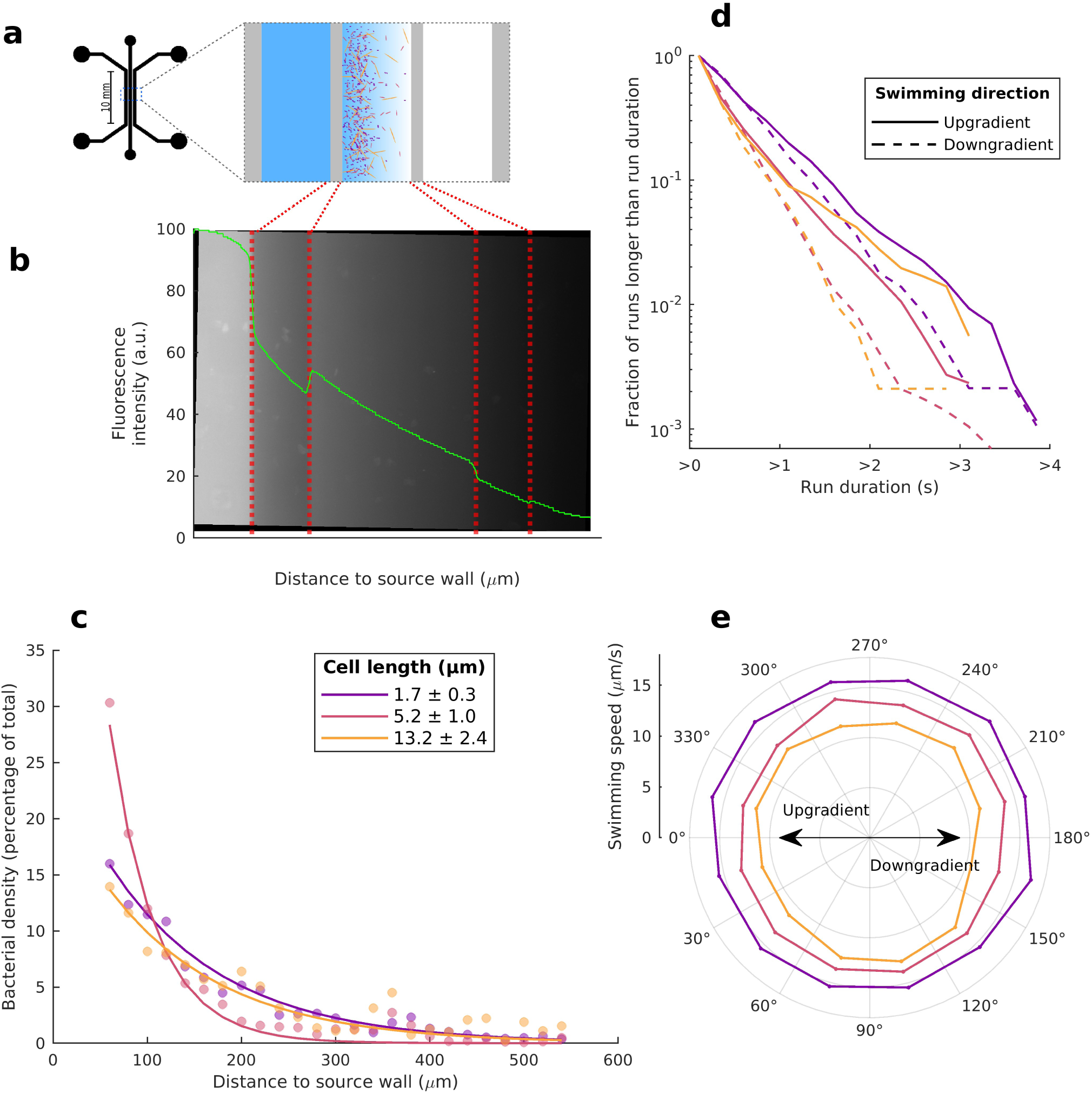
Chemotactic responses in example populations of E. coli of different cell lengths in a microfluidic device creating a linear gradient of α-methyl-aspartate. **a**) Schematic of the microfluidic device used for the chemotactic assays. **b**) Epifluorescence microscope image of a gradient of fluorescein in the microfluidic device showing the entire width of the central channel where bacteria are located, and partially the lateral channels were buffers with different amount of chemoattractant are flowing. Vertical dashed red lines mark the limits of the channels. Superimposed in green is the profile of average fluorescein concentration (in arbitrary units of fluorescence intensity). **c**) Bacterial cell densities across the width of the central channel for populations of E. coli of the three example cell lengths exposed to a linear gradient of α-methyl-aspartate (∇ *C* / *C̅* =1.2 mm^−1^). The shortest cells (1.7 ±0.3 µm, in purple) are untreated E. coli cells. The other two examples correspond to the average length of populations two hours (5.2±1µm, in red) and three hours (13.2±2.4 µm, in orange) after treatment with cephalexin. Circles are empirical measurements; continuous lines are the nonlinear least squares fits of the exponential. **d**) Inverse cumulative frequency of run times for cells swimming upgradient (continuous lines) and downgradient (dashed lines) with a maximum deviation of 45 degrees from the direction of the gradient. **e**) Polar plot of the average run speeds per mean direction of the runs.

### Chemotactic assays

Chemical gradients were generated by pumping in the outer channels of the device RMB with different concentrations of α-methyl-aspartate. In all experiments we kept the concentration of aspartate in the source channel at 100 µM, while changing the concentration at the sink channel to achieve different gradients. We performed experiments with three different sink concentrations (0, 25 and 50 µM) which, after accounting for the differences from theory quantified in the device calibration, resulted in cross-channel average relative concentration gradients of 1.20 mm^-1^, 0.72 mm^-1^ and 0.40 mm^-1^ respectively. Relative gradients are defined as 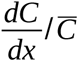, where *C* is the concentration of α-methyl-aspartate, *x* is the distance along the gradient and *C̅* is the average concentration over the whole width of the channel (see a list of all symbols used throughout the text in Table 1). These linear gradients were similar to those considered in previous experimental and theoretical studies of bacterial chemotaxis (e.g. Kalinin et al. 2009; Ahmed et al. 2010).

**Table 1:** Summary of the symbols used for each parameter and their definition.

|  | Symbol | Parameter | Definition | Units |
| --- | --- | --- | --- | --- |
| **Population-level parameters** | $C$ | Chemoattractant concentration | Environmental concentration of aspartate or α-methyl-aspartate | $\mu\text{M}$ |
| | $L$ | Chemotactic precision length-scale | Inverse of the exponential decay constant of bacteria distribution vs distance to nutrient source | $\mu\text{m}$ |
| | $t_{ss}$ | Time to steady state | Time it takes for IBM simulation to reach steady state, which is defined as the moment when the relative rate of change in $L$ is lower than 10% per hour | min |
| | $B$ | Normalized cell density | Density of bacteria per distance to source boundary, normalized to number of bacteria in the simulation | - |
| | $N$ | Normalized nutrient exposure | Population-averaged, time-integrated exposure to chemoattractant for the entire population in IBM simulations, normalized by integrated exposures for non-chemotactic populations distributed uniformly along the gradient | - |
| | $\mu$ | Bacterial random motility | Coefficient of diffusion of a bacterial population in the absence of chemical gradients, calculated fitting Taylor's equation (Taylor 1922) to tracks from IBM simulations of bacteria swimming in homogeneous conditions (Schuech and Menden-Deuer 2014) | $\mu\text{m}^2\text{s}^{-1}$ |
| | $v_c$ | Chemotactic speed | Mean speed at which bacteria move up a chemoattractant gradient | $\mu\text{m s}^{-1}$ |
| **Individual motility parameters** | $f_t$ | Translational friction coefficient | Translational friction coefficient along the long semi-axis of a prolate ellipsoid of revolution | $\text{kg s}^{-1}$ |
| | $f_r$ | Rotational friction coefficient | Rotational friction coefficient about the short semiaxes of a prolate ellipsoid of revolution | $\text{kg } \mu\text{m}^2\text{s}^{-1}$ |
| | $D_r$ | Rotational diffusivity | Rate of rotational diffusion about the short semiaxes of a prolate ellipsoid of revolution: $D_r = \frac{kT}{f_r}$ | $\text{rad}^2\text{s}^{-1}$ |
| | $\tau_r$ | Run time | Average duration of runs | s |
| | $\tau_t$ | Tumble time | Average duration of tumbles | s |
| | $TB$ | Tumble bias | Proportion of time spent tumbling: $TB = \frac{\tau_t}{\tau_t + \tau_r}$ | - |
| | $\alpha_p$ | Directional persistence parameter | Average of the cosine of the reorientation angles | - |
| | $v$ | Swimming speed | Average swimming speed during runs | $\mu\text{m s}^{-1}$ |

We imaged cells in a phase contrast inverted microscope (Zeiss AxioVert A1) equipped with a temperature-controlled stage (PE100-ZAL System, Linkam Scientific Instruments Ltd., Tadworth, UK) and using a 10X objective (0.55 NA), which gave a depth of field of 12.4 µm. Videos were recorded using a sCMOS camera (Hamamatsu ORCAFlash2.8, Hamamatsu City, Japan). The resulting lateral resolution was of 2.75 µm pixel^-1^, and the field of view was 524 µm by 598 µm, allowing the visualization of almost the entire width of the channel at once. Videos were taken at 30 fps which is fast enough to efficiently track *E. coli* (Guadayol et al. 2017).

In each experiment, several one-minute-long videos were taken at regular intervals starting 10 minutes after bacteria were seeded in the centre channel, giving ample time for the chemical linear gradient to re-establish (Ahmed et al. 2010), and ending 60 minutes after seeding. Motile bacteria were tracked using in-house MATLAB code described elsewhere (Guadayol et al. 2017) and published under an open-source license (Guadayol 2016). The code characterizes shapes and positions of individual cells and registers frequency, duration, speed and direction of runs, as well as frequency, duration and angle of reorientations.

A few minutes after injecting the bacterial culture into the central channel, cells visibly aggregated towards the outer channel with the highest chemoattractant concentration, following an exponential distribution. The inverse of the exponential decay constant, the “chemotactic precision length scale” (*L*), was used to parameterize chemotactic performance (Kalinin et al. 2009; Son et al. 2016). Precision length scales were estimated for each cell-length class and chemical gradient by fitting an exponential function to the distribution of bacteria across the centre channel using the Least Absolute Residual (LAR) robust curve fitting algorithm. Data points less than 50 µm from the walls were discarded because light diffracted by the agarose walls interfered with the particle detection algorithm. Other metrics of chemotactic response were considered but were ultimately disregarded because of large errors associated to their estimation in the experimental setup, and because of their sensitivity to the local gradient (*Supplementary material*, *Choice of response parameter*).

An occasional imperfect seal between the agarose and the PDMS resulted in shallower nutrient gradients and lower bacterial densities than expected. We identified experiments with deficient seals as those that met any of the following criteria: 1) residual flow in the centre channel was above 2 µm s^-1^ (i.e., ∼10% of the average speed for *E. coli*); 2) average number of cells detected was lower than 500; 3) distribution of cells was not monotonically decreasing away from the nutrient source, and 4) coefficient of determination *R^2^* of the fitted exponential model was <0.1. Overall, we recorded 126 videos, of which 89 were successful and used in subsequent analyses. All experimental data, including experiments not included in the final analyses, have been uploaded to a public data repository (Guadayol et al. 2024).

### IBM model

We developed an individual-based model (IBM) to gain a mechanistic insight into how chemotactic responses depend on cell length. The IBM was written in MATLAB code and published under GNU license (Guadayol 2023). The IBM simulates cells behaving independently (i.e. without interacting or communicating with other cells) in a 2D domain bound by parallel reflecting walls 600 µm apart, which is the width of the channel in the experimental microfluidic device. The dimensionality of the model not only replicates that of the microfluidic device, in which nutrient concentration is invariant in the vertical, but also of the motility parameters, which were estimated from 2D videos. The chemotactic behaviour of the cells in the IBM was simulated with a coarse-grained chemotactic pathway model described extensively elsewhere (Tu et al. 2008; Kalinin et al. 2009; Shimizu et al. 2010). See *Supplementary material*, *Chemotaxis signalling pathway model* for details on our implementation. Both the number of cells (10^5^) and the time step (0.1 s) were selected to ensure errors below 10% (*Supplementary material, Convergence tests*). Each simulation was randomly populated with cells of a given cell length. The number of cells remained constant throughout the simulation. Their “swimming speed” (*v*), “tumble bias” (*TB*, the proportion of time spent tumbling) and “directional persistence” (*α_p_*, defined here as the average of the cosine of the reorientation angles) were drawn from published empirical functional relationships of these parameters versus cell length for *E. coli* swimming in homogeneous conditions (Guadayol et al. 2017, *Supplementary material* Fig. S1). All parameters used in the IBM are listed in Table 1.

During a run, each cell was assumed to move at a constant run speed, whereas its swimming direction was affected by rotational Brownian motion. To account for this effect, after each time step, the run direction was changed by adding a normally distributed stochastic component with standard deviation 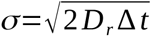, where *D_r_* is the size- and shape-dependent rotational diffusivity perpendicular to the long axis and `Δ`*t* is the time step of the simulation. Rotational diffusivity can be calculated as 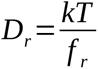, where *k* is the Boltzmann constant, *T* is the absolute temperature and *f*_r_ is the rotational friction coefficient about the short axes. We modelled bacteria as rigid prolate ellipsoids of revolution (Dusenbery 1998), and used well-known theoretical solutions for the friction coefficients (Perrin 1934).

The duration of both runs and tumbles was modulated stochastically from Poisson processes whose frequencies are the inverse of the expected durations and which generate the exponential distributions observed experimentally (Berg and Brown 1972). The expected duration of a tumble (*τ_t_*) changed only with cell length, whereas the expected duration of a run (*τ_r_*) depended also on environmental chemoattractant concentrations (see following subsections).

At the start of a simulation, cells were distributed uniformly and oriented randomly within a 600 by 600 µm square domain. A reflecting boundary condition was imposed at the two edges of the domain perpendicular to the nutrient gradient, analogous to the walls of the centre channel in the chemotactic device; our results were insensitive to this choice (*Supplementary material*, *Boundary behaviour, Fig. S2*). Cells bumping into one of the two edges parallel to the gradient were translated to the other boundary without a change in course or speed (i.e. a periodic boundary condition).

#### Phycosphere model

The phycosphere was simulated with the same IBM model on an annular 2D domain, in which the inner circle approximated the wall of a spherical phytoplankton cell of radius 25 µm. The outer boundary was set to a radius of 500 µm, corresponding to half the average distance between neighbouring microalgae at a concentration of 10^3^ cells·mL^-1^. Bacteria encountering a boundary bounced away conserving the same angle of incidence and speed. A 2D domain was necessary to match the dimensionality of the measurements of size-dependent motility parameters (*Supplementary material* Fig. S1), which were obtained from 2D videos.

The chemoattractant concentration gradient around a spherical phytoplankton cell at steady state, assuming background concentrations to be 0, follows *C* (*r*)=*C*_0_ *r*_0_ *r*^−1^ (e.g. Jackson 1987), where *r* is the radial distance from the centre of phytoplankton cell, *r*_0_ is the radius of the cell, and *C*_0_ is the concentration of the chemoattractant at the cell wall. *C*_0_ was set to be constant (corresponding to a situation where the cell is releasing the chemoattractant at a constant rate) with an arbitrary value of 10 µM that ensured concentrations of aspartate within the sensitivity range of *E. coli*, which is a poor chemotacter in comparison with free-living planktonic bacteria (Stocker 2011). In this scenario, at steady state the shape of the chemical gradient is independent of molecular diffusivity.

Phycosphere simulations were run for 10 minutes to match reported temporal spans of formation and dissipation of bacteria aggregates around nutrient hotspots (Blackburn et al. 1998; Smriga et al. 2016).

## Results

### Experimental results

As cells elongated, their motility pattern changed: swimming speeds decreased, tumble angles were narrower, runs shorter and tumbles longer (Fig. 2, *Supplementary material* Fig. S1). We detected chemotactic responses in populations of *E. coli* of all cell lengths. As is commonly observed in *E. coli*, these responses were caused by a directional bias in run times rather than in swimming speed (Fig. 1c, d). After a few minutes, cells distributed along the chemical gradient following an exponential relationship in all experiments (Fig. 1b) of the form *B* (*x*)=*B* ( 0 )*e*^−*x*/*L*^, where *B*(*x*) is the cell density at distance *x* from the wall closest to the most concentrated outer channel, and *L* is the chemotactic precision length-scale that parametrizes how tightly cells aggregate near the source of chemoattractants (Son et al. 2016). The observed exponential distribution is the steady-state solution of the Keller-Segel’s bacterial transport model for a linear chemical concentration profile (Kalinin et al. 2009; Son et al. 2016). In this solution, at steady state *L*=*μ* / *v_c_*, where *µ* is the random motility coefficient that parametrizes the diffusivity of bacteria in the absence of chemical gradients, and *v_c_* is the chemotactic drift velocity (i.e. the average speed at which bacteria swim towards the source). Thus, *L* can be interpreted as the ratio of bacterial diffusivity to chemotactic ability. The shorter *L* is, the better bacteria are at exploiting a nutrient source.

**Figure 2:**
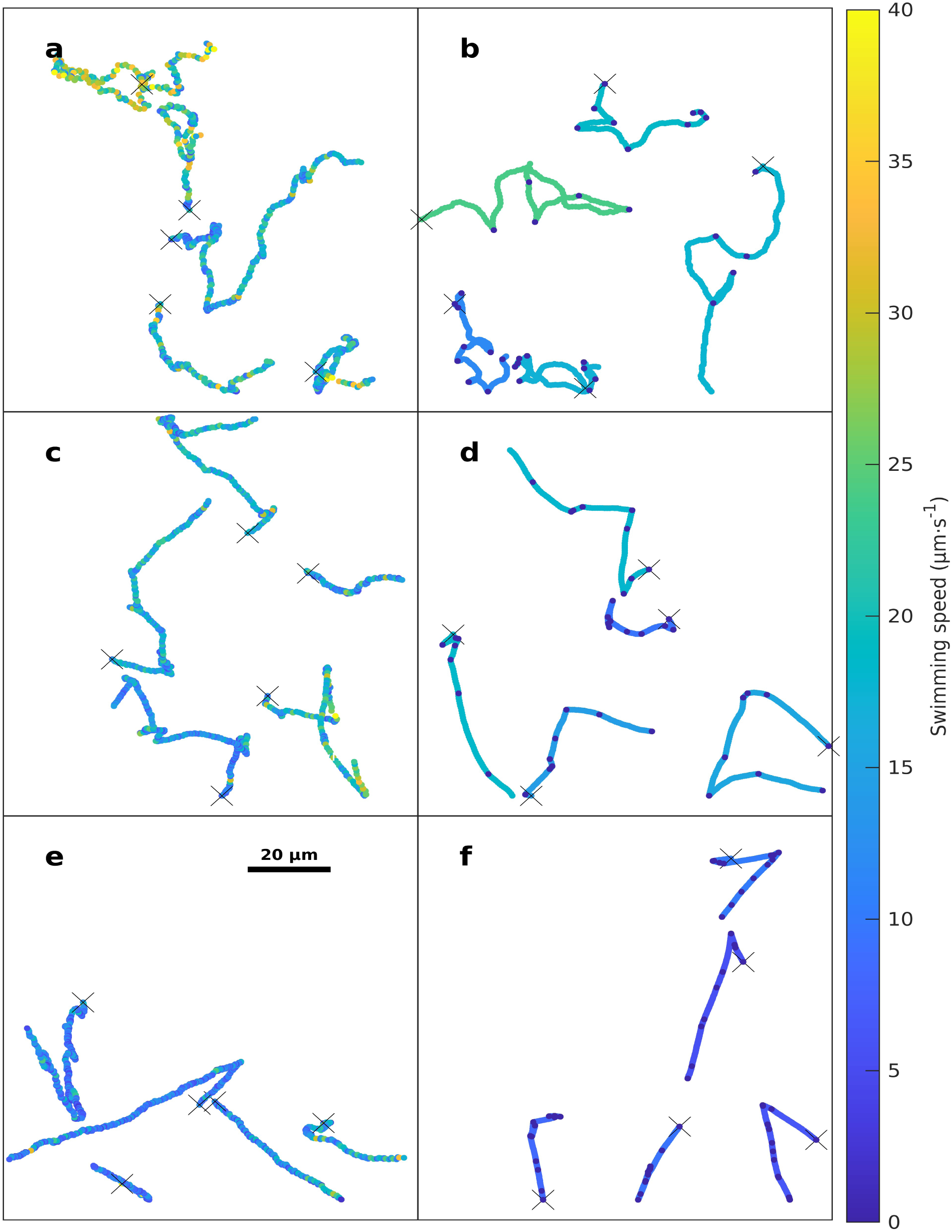
Example trajectories of cells of different length. **a**, **c** and **e** are randomly selected experimental trajectories of cells exposed to a linear gradient of α-methyl-aspartate. **b**, **d** and **f** are randomly selected trajectories from the corresponding IBM simulations. **a** and **b** show trajectories of cells 1,7 ±0.3 µm long (i.e. untreated E. coli cells); **c** and **d** show trajectories of 5.2±1µm long cells (i.e. ∼two hours after cephalexin treatment); **e** and **f** show trajectories of 13.2±2.4 µm long cells (i.e. ∼three hours after cephalexin treatment).

As previously observed (Kalinin et al. 2009), *L* decreased with increasing gradient steepness (Fig. 3). Regardless of the steepness, the functional response of *L* to cell length was U-shaped, with minimum *L* occurring for cells 2.5 to 5 µm long. This supports our hypothesis that some trade-off for optimal chemotaxis is operating in *E*. *coli*. However, our experimental system alone does not allow identifying which mechanisms are driving the observed pattern for two reasons. First, *L* is an emerging population parameter that ultimately depends on at least four shape-dependent individual-based motility parameters. These parameters are rotational friction coefficient *f_r_*, swimming speed *v*, tumble bias *TB* (that is the proportion of time spent tumbling), and directional persistence *α_p_* (defined here as the average of the cosine of the reorientation angles). All these parameters have non-linear functional relationships with cell length (*Supplementary material* Fig. S1, Guadayol et al. 2017) and therefore their individual effect cannot be isolated. Thus, the observed pattern has multiple possible non-mutually exclusive explanations. For example, high *L* could be equally explained by conditions that increase *µ*, such as high *v* or *α_p_*, or by conditions that decrease *v_c_*, such as low *f_r_*. A summary of these possible explanations based on current theories is given in the *Supplementary material*, *Theoretical background* and Table S1.

**Figure 3:**
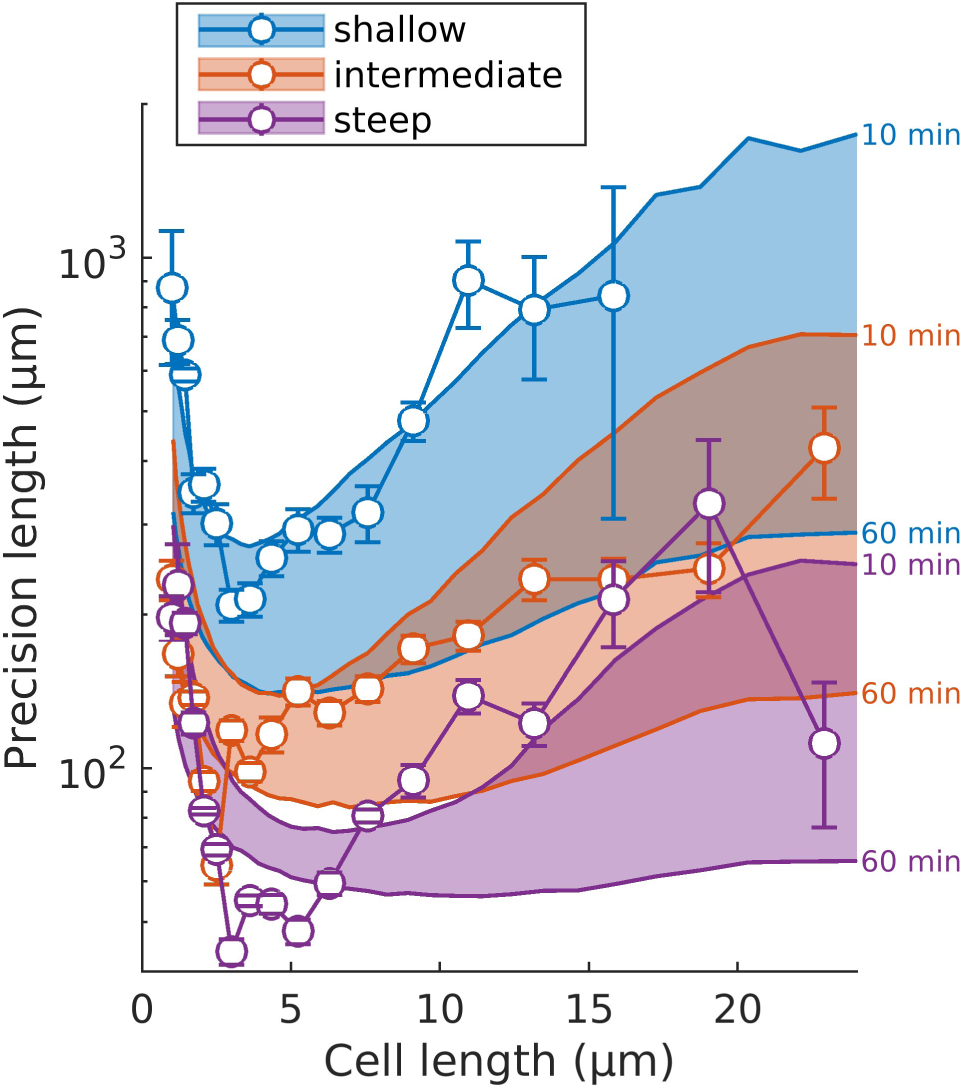
Experimental and simulated precision length scales vs. cell length of E. coli populations exposed to linear gradients of α-methyl-aspartate of different steepness for a period between 10 and 60 minutes. Colours represent the three gradients used in the experiments, blue being the shallowest gradient, red the intermediate and purple the steepest. Data points are experimental results, with error bars showing the standard errors of the exponential fits. Coloured areas show the IBM model outputs between 10 (upper limits) and 60 (lower limits) minutes of simulation time.

The second reason that prevents identifying underlying mechanisms is that we manipulated cells into adopting potentially unnatural shapes, which could induce experimental artefacts. For example, flagella could become increasingly desynchronized as cells elongate, because of the longer distance that chemical signals need to diffuse and because of the difficulty of forming a single bundle when flagella are far apart (Lee et al. 2021). This might lead to changes in *TB* that may partly explain the increase in *L* for the longest cells. However, such experimental artefacts do not necessarily reflect the biophysical constraints on chemotaxis over evolutionary time and make it difficult to generalize results.

In order to tease apart which parameters are driving the observed pattern, we developed an individual-based model (IBM) that simulates a population of chemotactic bacteria of a given cell length swimming along a linear gradient of aspartate. Model simulations between 10 and 60 minutes (corresponding to the experimental time span) show good fits with the experimental data for the three chemoattractant gradients explored (Fig. 3).

### IBM dynamics

In all IBM simulations, *L* monotonically decreased over time, approaching a horizontal asymptote (*Supplementary material* Fig. S3). We defined an operational steady state as the point at which the relative rate of change in *L* was lower than 10% per hour. The time *t_ss_* it took for simulations to reach such steady state (grey circles in Fig. 4b) replicated the U-shaped relation with cell length and ranged between 31 and 107 minutes. This indicates that experiments were not long enough for populations to achieve steady state distributions, particularly for cells longer than 10 µm. Allowing simulations to proceed until steady state revealed a trend different to what was observed experimentally: *L* decreased almost monotonically with cell length (Fig. 4a) instead of showing a local minimum at intermediate cell lengths (Fig. 3). This divergence between long term simulations and experiments and simulations shorter than 60 minutes highlights the transient nature of the measurements. A more ecologically relevant parameter, the population-averaged time-integrated exposure to nutrients *Ν*, shows the relative advantage of cells of intermediate length over the longest cells to extend for simulations as long as 2 hours (Figs. 4c, 5a). This advantage, however, diminishes as simulation time increases (Fig. 5a).

**Figure 4:**
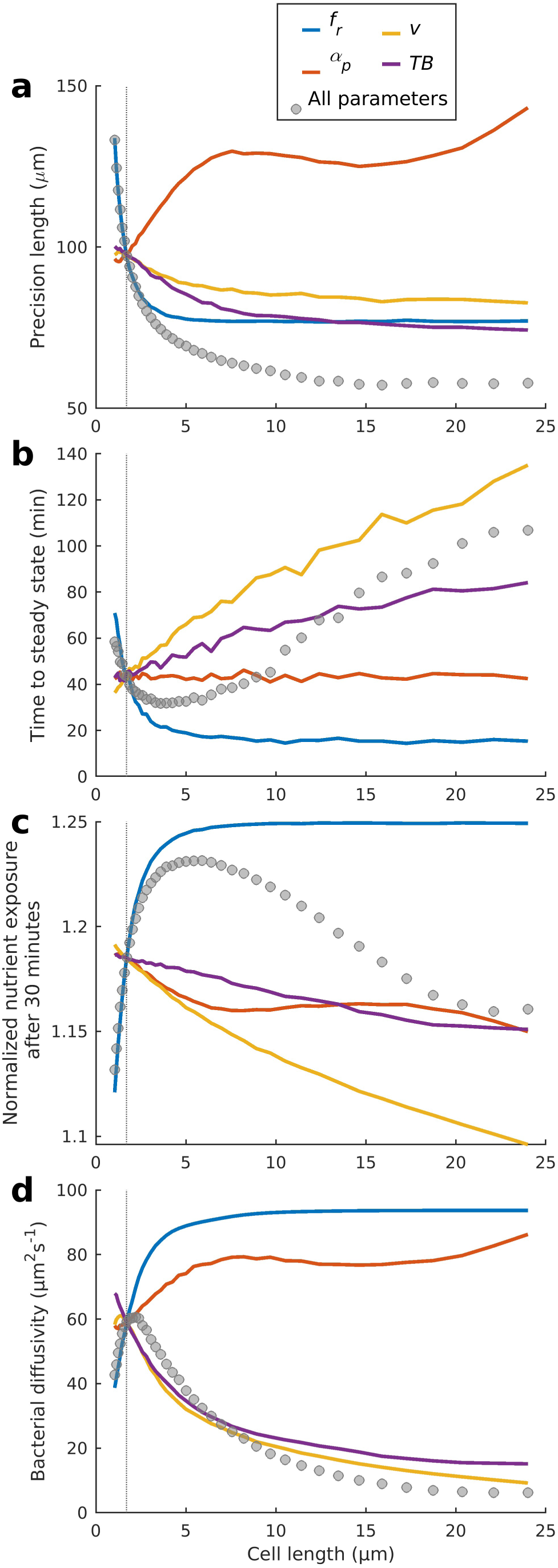
Sensitivity analyses at steady state of the IBM. **a)** Precision length scales L at steady state; **b)** time t_ss_ it takes for simulations to reach steady state. **c)** Normalized exposures to α-methyl-aspartate N after 30 minutes of simulation time. **d**) Bacterial diffusivities in homogeneous conditions. Grey circles represent simulations where all parameters changed with cell length according to the empirical relationships for cephalexin-treated E. coli shown in Supplementary material Fig. S1. Coloured lines show the results of the one-at-a-time sensitivity analysis, where one parameter varies with cell length while the rest are fixed to the values corresponding to normal-size wild-type E. coli.

**Figure 5:**
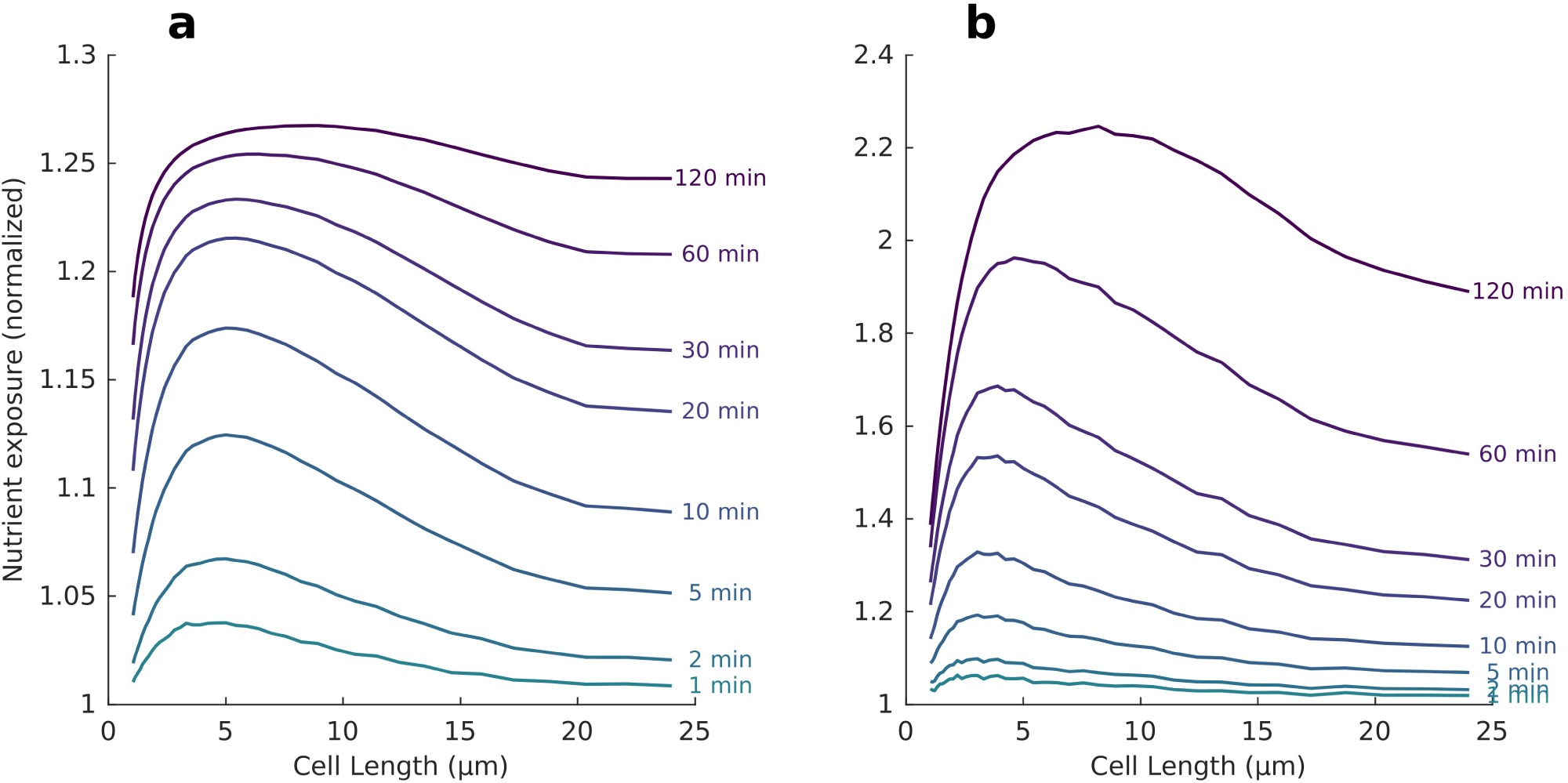
Time-integrated nutrient exposure per cell vs cell length in simulations of A) a linear gradient of aspartate, and B) a phycosphere of 25 µm radius and a concentration of Asp at the surface of the microalga of 10 µM. Values are normalized by the average nutrient exposure of a non-chemotactic population uniformly distributed across simulation space.

The random motility coefficient (*µ*), which parametrizes the diffusivity of bacteria in the absence of gradients (and is therefore a proxy for exploration), shows also a maximum at intermediate lengths (Fig. 4d), although this occurs at shorter cells and is narrower than for nutrient exposure.

### Sensitivity analysis

To understand the individual role of each motility parameter in the emerging pattern, we performed a sensitivity analysis of the model (Fig. 4) using a one-at-a-time (OAT) approach: we ran a series of simulations to steady state in which we changed each one of the relevant shape-dependent parameters (*f_r_*, *v*, *TB*, *α_p_*) whilst keeping the other parameters constant at the empirical average values for a wild-type cell 1.7 µm long and 0.75 µm wide at a temperature of 30°C swimming in a homogeneous environment (*f_r_*=0.4·10^-20^ kg m^2^ s^-2^, *v*=18 µm s^-1^, *TB*=0, *α_p_*=0, *Supplementary material* Fig. S1, Guadayol et al. 2017). Because tumble time *τ_t_* is insensitive to changes in chemoattractant in the chemotactic pathway model, responses to *TB* mostly reflect responses to run time *τ_r_*.

This analysis showed that, at steady state, changes in *v*, *TB* and, most noticeably, *f_r_* drove decreases in *L* as cells elongated (Fig. 4a). On the other hand, *α_p_* drove increases in *L* that may moderate the decay in *L* as cells elongate. The operating trade-offs are more clearly revealed in the steady state time (*t_ss_*, Fig. 4b) and, ultimately, in the integrated exposure to nutrients (*N*, Fig. 4c) and random motility (*µ*, Fig. 4d): slow performances of short cells are explained by *f_r_*, whereas those of long cells are driven by *v* and *TB*. Steady state time remained insensitive to changes in *α_p_*.

### Chemotaxis in the phycosphere

One of the most important hotspots of microbial activity in aquatic systems is the phycosphere, the volume influenced by the metabolic activity of a phytoplankton cell that shows a chemical composition distinct of background seawater and where many of the interactions with other microorganisms occur (Seymour et al. 2017; Raina et al. 2022). To challenge our results for cell length dependent performance against an ecologically relevant scenario, we further used our model to explore which shape may be optimal for chemotactic bacteria seeking phycospheres.

We considered the diffusion of a chemoattractant from a 25 µm radius phytoplankton cell with a constant rate of release and assumed steady state following previous models (Stocker 2011). In this scenario, concentration does not decrease linearly as in our chemotactic device but as a function of 1/*r*, where *r* is the radial distance to the centre of the phytoplankton cell. We then ran IBM simulations within a 2D plane to model the clustering of bacteria around phytoplankton. The results show bacterial distributions consistently peaking some micrometres away from the phytoplankton cell surface (Fig. 6b). This is a behaviour commonly observed in IBMs of chemotaxis and is called the “volcano effect”(e.g. Bray et al. 2007; Simons and Milewski 2011). It results from bacteria swimming past the point of maximum concentration of attractant because it takes some time for the cell to detect a decrease in chemoattractant concentration and initiate a tumble. This reaction time is often parameterized by the latency time, which is a measure of the time needed for signals generated by external stimuli to be processed through the complete signal transduction pathway. A sensitivity analysis indicated that although the magnitude of this local peak was sensitive to changes in all motility parameters, its location was most sensitive to swimming speed (*Supplementary material* Fig. S4) because, given chemotactic response time, faster swimming translates into longer reaction distances. Thus, for example, in our modelling framework the distribution of a population of bacteria swimming at 20 µm s^-1^ peaked 40 µm away from the phytoplankton cell wall, whereas that of a population of bacteria swimming at 100 µm s^-1^ peaked 200 µm away.

**Figure 6:**
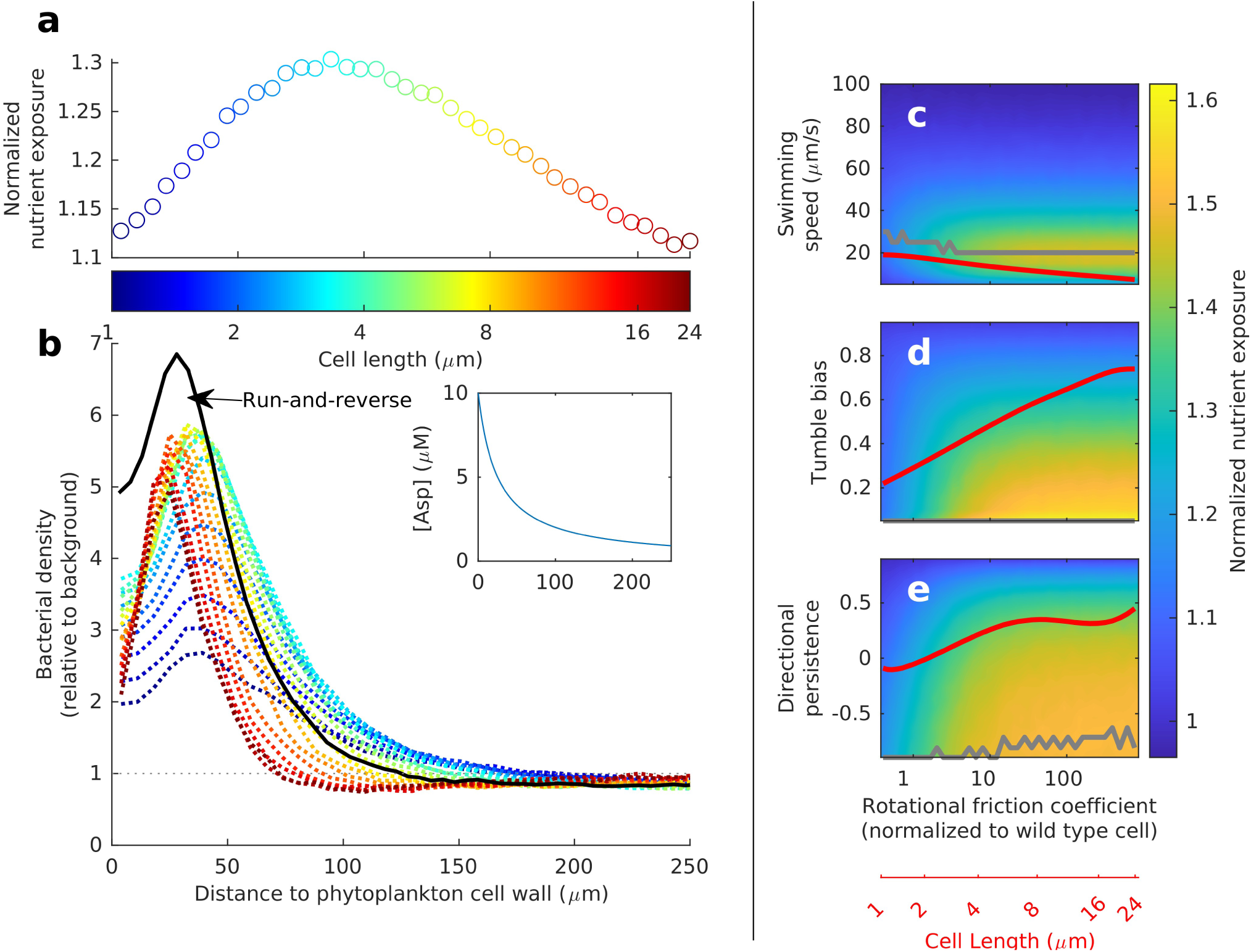
Modelled chemotactic performances in a phycosphere. **a**) Normalized integrated nutrient exposure (N) after 10 minutes of simulation time, against cell length for simulated populations of E. coli around a spherical phytoplankton cell of radius 25 µm releasing Asp at a constant rate. **b**) bacterial densities normalized by background densities vs distance to phytoplankton cell wall. Colour-coded dashed lines represent populations of increasing cell length. The black line represents simulation for a population of wild-type E. coli cells displaying a run-and-reverse motility pattern, common in marine planktonic motile bacteria. Run-and-reverse pattern was modelled enforcing a constant tumble angle of 180 degrees for all reorientations. Inset shows the distribution of the chemoattractant Asp used in the simulations. **c-e**) integrated nutrient exposure N as a function of rotational friction coefficient f_r_ and swimming speed v (**c**), tumble bias TB (**d**), and directional persistence α_p_ (**e**). Although v, TB, and α_p_ can all be treated as independent of f_r_ in our simulations, the red lines show the observed dependences derived from experimental work with E. coli (Guadayol et al. 2017). Grey line shows the maximum nutrient exposures per rotational diffusion within each two-parameter space.

The time-integrated exposure to nutrients *N* is again a better measure of the ecological relevance of these results. As in the linear gradient case, *N* increases rapidly with cell elongation up to a length of around 3.3 µm and then decreases more gradually (Fig. 5a). To understand this trend, we explored the 2-D spaces defined by *f_r_* on one axis and *v*, *TB*, and *α_p_* on the other (Fig. 6c-e). This exercise shows that as cells elongate, *N* is maximized by low *TB* (i.e. long runs), an *α_p_* close to −1 (characteristic of a run-and-reverse pattern), and a speed *v* of around 20 µm s^-1^. Slow swimming leads to slow aggregation around the phytoplankter, but swimming too fast amplifies the volcano effect (*Supplementary material* Fig. S4).

## Discussion

### Physical constraints and biological trade-offs

The time-integrated exposure to chemoeffectors *N*, which we consider a measure of fitness of a chemotactic strategy, depends on two factors: the precision and the speed of chemotaxis. All shape-dependent motility parameters affect these two factors in different ways (Fig. 4). Chemotactic precision, which we parameterize with the chemotactic precision length-scale *L*, improves sharply with increasing *f_r_*, and to a lesser extent, with decreasing *v* and increasing *TB*. The result is that, at steady state, precision improves as cells elongate. On the other hand, the speed of the chemotactic process, which is inversely related to the time *t_ss_* to reach steady state, is positively related to *v* and *f_r_* and negatively related to *TB*. The result is that chemotaxis is fastest for *E. coli* cells of 3 to 7 µm in length. Thus, the U-shaped response of *L* to elongation that we observed empirically likely resulted from shape-dependent motility parameters acting oppositely on the precision versus speed of the chemotactic process. Thus, alongside the response of random motility (*µ*) in the absence of gradients (Fig. 4d), our results reflect the well-known trade-off between exploitation and exploration operating in chemotactic bacteria in aquatic systems (Clark and Grant 2005; Altindal et al. 2011).

The general trends observed in *L* at steady state in response to changes in *f_r_*, *v,* and *α_p_* conform to predictions from theoretical models. For example, the response of *L* to elongation-driven changes in *f_r_* (Fig. 4a) is consistent with the monotonic improvement in gradient detection predicted by a previous study based purely on length-dependent changes in translational and rotational drag in ellipsoids of revolution (Dusenbery 1998). The decrease in *L* in response to decreases in *v* as cells elongate is consistent with the Keller-Segel model, which predicts a positive, linear relationship between these parameters (Rivero et al. 1989). Finally, the increase in *L* with increasing *α_p_* as cells elongate is consistent with predictions from a previous study (Lovely and Dahlquist 1975) that established a positive relation between *α_p_* and the random motility parameter *µ* (and hence, also *L*).

The apparently small effect that *α_p_* has on *L* and *t_ss_* (Fig. 4a,b) is inconsistent with a strong positive relationship between *α_p_* and chemotactic velocity *v_c_* predicted by a previous analytical model (Locsei 2007). This strong relationship should induce decreases in precision length (since *L*=*µ*/*v_c_*) and in *t_ss_* as cells elongate. Instead, *L* increases and *t_ss_* remains invariant, suggesting that the effect of *α_p_* on *µ* is overpowering its expected effect on *v_c_*. Alternatively, the marginal significance of *α_p_* in our results may also be reflecting its poor performance as a descriptor of reorientation behaviour, at least in peritrichous bacteria such as *E. coli*. Motility patterns are defined by the mode of reorientation between runs. The changes in motility pattern imposed by elongation in *E. coli* are poorly characterized by *α_p_* because it does not discriminate between 180° reorientations and true reversals (where a leading pole in the previous run becomes the trailing pole, as the bundle is reformed in the opposite pole). Thus, for example, *α_p_*=0 could equally result from a truly uniform distribution of tumble angles and from a pattern of angles in which half are 0° and half are 180°. In any case, the relationship between *α_p_* and *L* needs to be further explored.

### Ecological and evolutionary implications

The optimal values for chemotaxis in our simulations may not be directly ecologically representative because the IBM is based on *E. coli*’s chemosensory circuit for aspartate and uses species-specific motility patterns and physiological responses to elongation. We therefore expect our results to be somewhat dependent on species-specific sensitivities to different chemoattractants and to the magnitude and shape of the chemical gradient, as well as the more generic aspects of the geometry and boundary conditions of the model domain, and the simulation time. However, we suggest that our results still reflect underlying trade-offs and evolutionary pressures acting on bacterial morphology.

Our results indicate that an optimal value for cell length arises from the well-known trade-off between the need for rapidly tracking nutrient patches (exploration) and the need for remaining close to them (exploitation) (Clark and Grant 2005; Altindal et al. 2011). Long cells will eventually maintain a tighter distribution near a nutrient source whereas short cells will track gradients faster. Thus, the optimal cell length of bacteria realized in nature will likely depend on the particulars of their environment (Stocker and Seymour 2012). In environments where gradients are long and stable over time (relative to microbial scales), such as at the sediment-water interface, we may expect selective pressure for remaining close to the nutrient source and therefore for increasing directional control by increasing *f_r_* and *α_p_*. This is consistent with observations of extremely long and large bacteria near and within sediments (Fenchel 1994; Guerrero et al. 1999; Jørgensen 2006). In the planktonic environment, with its patchy, dynamic and ephemeral nutrient landscape, speed is likely more important (Xie and Wu 2014). This may partly explain why the median aspect ratio of planktonic bacteria is relatively small (around three:Dusenbery 1998), and why flagellated bacteria in oligotrophic environments tend to swim faster (Mitchell et al. 1995). This argument, of course, does not reflect the fact that tasks and costs other than chemotaxis may be important for bacterial fitness (Young 2006; Schuech et al. 2019), nor their interaction with flow (Luchsinger et al. 1999; Taylor and Stocker 2012).

This need for speed in the planktonic environment may be moderated, as our simulations show, by the occurrence of a volcano effect that penalizes very fast swimmers (Fig 6, *Supplementary material* Fig. S4). The volcano effect results from cells overshooting the phycosphere as it takes some time (the “latency time”) for the bacterial chemotactic pathway to perceive a decrease in chemoattractant concentration and induce reorientation (Bray et al. 2007; Simons and Milewski 2011). Thus, the importance of the volcano effect depends on three factors: the shape and length of the chemoattractant gradient (which will themselves depend on the size of the phytoplankton cell), the latency time, and the swimming speed. The slower the cells are at reacting to a change in concentration and the faster they swim, the further from the maximum concentration will their population distribution peak. In our simulations this translates into optimal nutrient exposures at swimming speeds remarkably close to the average value for wild-type *E. coli* (Fig. 6c). In general, the latency time will add a constraint on the maximum swimming speed that is ecologically viable in a given planktonic system.

If the volcano effect proves to be an ecologically relevant mechanism, there are several factors that could be reducing its impact in planktonic bacteria in aquatic systems. First, their latency times are thought to be much faster (Stocker 2011) than those of *E. coli* (∼0.2 s Block et al. 1982). Second, speed-dependent responses such as chemokinesis or changes in motility pattern (Son et al. 2016), which we have not considered here, may help bacteria remain longer around nutrient patches even when their swimming speeds are high. Third, the adoption of swimming patterns with a reversal phase and long runs, which are prevalent among marine bacteria (Johansen et al. 2002) can also maximize nutrient exposure around a phycosphere (Fig. 6). And finally, the existence of a viscous layer of exopolymers around a phytoplankton cell may minimize the volcano effect (Guadayol et al. 2021).

Otherwise, our simulations of the phycosphere depict an optimal strategy that is largely consistent with observations of marine bacteria. The simulations predict that elongated cells with a run-and-reverse pattern (i.e. *α_p_*→-1) and long runs (i.e. *TB*→0) will be most efficient in harvesting nutrients at timescales of tens of minutes (Fig. 6). Among marine bacterial populations, the median cell aspect ratio is close to 3 (Dusenbery 1998), the prevalent motility pattern is run-and-reverse (Johansen et al. 2002) or has some sort of reversal phase (Xie et al. 2011), and the most common type of flagellation is a single polar flagellum (Grognot and Taute 2021), which favours long runs at the cost of increasing chemotactic response times (Sneddon et al. 2012).

We have focused on individual motility parameters (rotational friction, run speed, tumble and run times, directional persistence) that are dependent on the shapes of the organisms in our experimental system, and that therefore may be expected to change in natural populations displaying morphological variability. However, cell shape is not the only factor sensitive to selective pressures that can influence these motility parameters. For example, high *f_r_* can also be achieved by means of long flagella, which can act as stabilizers of the running trajectory (Mitchell 2002; Schuech et al. 2019). Such stabilization would allow cells to be smaller while remaining chemotactically efficient. Similarly, changes in *v* or *α_p_* can be achieved by means other than elongation, for example by alterations in the number or positions of flagella (Grognot and Taute 2021). Nevertheless, given the morphological diversity often seen in natural populations, and the ease with which some cell populations appear to change shape, and in particular aspect ratio, varying length may well be a response to an ever-changing nutrient landscape.

Chemotaxis is but one of several activities that may exert selective pressures on bacterial shape in general and cell length in particular (Young 2006). Some of these pressures may in fact act in opposition to the chemotactic mechanisms we reveal here. For example, specific nutrient uptake is highest in very small and filamentous cells. Protozoan grazers predate most efficiently upon bacteria of intermediate sizes and thus may select for small cells and filaments (Pernthaler 2005). And finally, passive dispersal by Brownian motion is most efficient for small spherical cells (Dusenbery 1998). Regardless, the fact that slightly elongated rods are the prevalent morphology among marine motile bacterioplankton hints at the important role that motility and chemotaxis play in shaping natural populations of aquatic bacteria.

In summary, our work dissects the main physical mechanisms that explain the prevalence of slightly elongated forms among chemotactic bacterioplankton and showcases the interplay between them. Both in a linear gradient replicating the long, stable gradients near static boundaries, and in a non-linear gradient replicating a phycosphere in a much more dynamic planktonic setting, we reveal the existence of an optimal cell length for bacterial chemotaxis and show how it arises from a tradeoff between the needs for exploration and exploitation.

## Acknowledgements

We thank H. Fu for comments on previous versions of this manuscript. This study was funded by Leverhulme Trust project RLA RL-2012-022, “Form and function in a microbial world”, granted to S.H., and by a Gordon and Betty Moore Foundation Marine Microbiology Initiative award, grant 6852 to S.H. and O.G. O.G. was supported by a María Zambrano grant from the University of the Balearic Islands. No conflicts of interest are declared.

## Data availability statement

The dataset of videos generated and analysed during the current study is not publicly available due to storage restrictions but is available from the corresponding author on reasonable request. Bacterial trajectories and cell lengths extracted from the videos are published in a public repository (DOI: 10.6084/m9.figshare.25452355), alongside the details of the experimental conditions needed to reproduce all analyses and figures. The code developed and used in this study is available under an open-source license at Zenodo.org (DOI: 10.5281/zenodo.3452818; DOI: 10.5281/zenodo.8360667).

