## Supplementary material for "Optimal cell length for exploration and exploitation in chemotactic planktonic bacteria"

**Affiliations**:

### Material and Methods

#### Microfluidic devices

To manufacture the device, an SU-8 model of the device was first fabricated using standard photolithography techniques by microLiquid s.l. (Arrasate, Spain). A PDMS mould of the SU-8 was then cast and used repeatedly to produce the agarose devices. Each agarose device was manufactured before the experiment by pouring RMB with 3% (w/v) agarose previously heated on a microwave oven onto the PDMS mould. The agarose was pressed down with a glass slide to ensure a flat surface and a good seal in later steps. It was then allowed to gel at room temperature before peeling it off and mounting it facing up on a glass slide. A PDMS slab with holes for the inlets and outlets was placed on top of the agarose. To generate the gradient, motility buffers with different concentrations of the non-metabolising amino acid α-methyl-aspartate were continuously pumped through the outer channels at a rate of 5μL $min^{-1}$ using a PHD Ultra syringe pump (Harvard Apparatus, Holliston, Massachusetts, U.S.). Buffers were pumped in withdrawal mode as negative pressure improves sealing and helps preventing leaks without the need to clamp the device (Ahmed et al. 2010). After 10 minutes of starting the flow, sufficient for a steady linear gradient of α-methyl-aspartate to be established in the quiescent centre channel, this was seeded with the bacterial population of interest. After seeding, the chemical gradient quickly re-established itself in the centre channel as it was already formed in the underlying agarose floor (Ahmed et al. 2010). Bacteria were then monitored as they swam and aggregated towards the source channel.

We calibrated the microfluidic device by pumping a 100 μM fluorescein solution at 5 μL $min^{-1}$ through the source outer channel and ultrapure water through the sink channel at the same rate. We then measured the decay in fluorescence across the channel with a Nikon Eclipse 80i epifluorescence microscope fitted with a FITC filter (Fig. 1b). Images were taken every 10 seconds to evaluate the time it took for the gradient to establish and stabilize. While some nonlinearity was present, profiles were well approximated as linear (Fig. 1b). The slopes of the linear gradients generated in our setup were 60% of that predicted by the diffusion equation given the geometry of the device and assuming constant diffusivities throughout the device and no leakages (Ahmed et al. 2010). Such inconsistencies have been seen before and are believed to be caused by a steeper decay profile in the agarose ridges flanking the central channel (Ahmed et al. 2010).

#### Choice of response parameters

In choosing the parameter to evaluate chemotactic responses we considered several candidates, including parameters that directly capture chemotactic behaviour, such as chemotactic speed $v_c$ or chemotactic sensitivity $\chi_c$, parameters describing the distribution of populations across a chemical gradient, such as precision length $L$ or centre of mass of the chemotactic population, and ecologically relevant parameters such as integrated exposure to nutrients. Ideally, the diagnostic parameter should be independent of the experimental system (i.e. of the geometry and steepness of the gradient), so that it would be easily comparable to other results. In this regard, $v_c$ (usually normalized to swimming speed $v$) and $\chi_c$ are clearly preferable as they parameterize intrinsic responses of the organism to a specific chemoeffector. However, our preliminary analyses showed these parameters to be very sensitive to the local gradient (see for example Fig. S3), and therefore proved to be an unreliable metric in our experimental system.

Population-level parameters that describe the distribution of cells across the chemical gradient are time dependent, but in a closed system, such as the microfluidic device used in our experimental work, values converge (Fig. S3). We chose *L* because 1) it has a solid theoretical basis, as it is the solution of the Keller-Segel model for a linear gradient 2) it is robust even when number of cells are low and 3) it has been used in a number of previous studies with a similar experimental setup.

To highlight the ecological significance of our results, we also report the time-integrated, population-averaged exposure to nutrients *N* from model simulations. This parameter, unlike *L*, does not converge to a single value with time but it increases unbounded (Fig. S3), and thus results are reported for specific times of experimental or ecological significance.

#### Motility parameters' responses to elongation

Except for rotational friction coefficients, for which the cells were modelled as ellipsoids of revolution  (Perrin 1934), the motility parameter values for any given individual cell in a simulation were drawn randomly from empirical probability density functions (PDFs) whose parameters depend on cell length. The functional relationships between the PDF parameters and cell length were established from published observations of cephalexin-treated *E. coli* swimming in a homogeneous motility buffer (Guadayol et al. 2017) and are shown in Fig. S1. Briefly, for each motility parameter, experimental data were allocated into logarithmic bins of cell length. The bins were logarithmic to ensure approximately even samples sizes across bins and a better resolution at short cell lengths, where motility parameters show the steepest changes. Then, appropriate PDFs were fitted to each cell length class. Lognormal PDFs were fit to run speeds whereas exponential PDFs were fit to run and tumble times, which arise from Poisson processes (Berg and Brown 1972). Finally, to reduce experimental noise, the resulting empirical relationships were smoothed by fitting spline functions using the SLMtools MATLAB toolbox (D'Errico 2009).

Tumble angles were randomly generated for each individual reorientation event in each simulation. However, in this case the derivation of a smoothed empirical relation between the PDF of the tumble angles and cell length was not direct because the shape of the PDF changes as cells elongate. Thus, a polynomial surface was fitted directly to the log-transformed frequency of events plotted vs the tumble angles and the logarithm of cell length, using a bisquare linear least-square fitting algorithm (MATLAB® function "fit").

#### Model behaviour at walls

The behaviour of cells when hitting the walls of the chemotactic device is not well described and is a potential source of discrepancy between the model and the data (Kalinin et al. 2009). To evaluate the significance of this uncertainty, we simulated five different plausible behaviours before choosing one to run the simulations. In the first behaviour (“Bounce”), the boundary was reflective, that is, cells bounced against the wall and kept running with the same speed and the same angle with respect to the normal direction to the wall, but inverse direction. In the second behaviour (“Stick for 1 s”), cells stuck to the wall for 1 second before starting a new run in a random direction. This was the behaviour used by Kalinin (Kalinin et al. 2009). The third one (“Stick for 0.1s”) was similar, but cells were stuck to the wall for the average duration of a tumble. In the fourth scenario (“Tumble”), cells started a tumble immediately upon hitting the wall. Finally, in the last scenario (“Push against boundary”), cells tried to keep running in the same direction as they came, “pushing” against the wall until a tumble was started or Brownian motion led to a directional change away from the wall.

The first four scenarios resulted in similar bacterial distributions even near the walls. Any differences among them were confined mostly to the first five micrometres near the chemoattractant source (Fig. S2). As the exponential fitting was only performed in the range 50 to 550 µm for coherence with the analyses of experimental data, these differences did not translate into differences in the precision lengths except for the last behaviour, which rendered the results least similar to the experimental observations. In our simulations we used the first behaviour (“Bounce”).

#### Convergence tests

We tested convergence of the model with respect to key numerical parameters by first running a series of simulations with an increasing number of cells (500 to 128000). For each number of cells, 40 simulations were run to assess accuracy and precision of the estimates. We ran this analysis with three different sets of chemotaxis parameters that yielded the extreme values of $L$ and steady state times $t_{ss}$ encountered in this study (Fig. 3). No significant differences in mean values were detected for any of the three sets of chemotaxis parameters for number of cells > 4000 individuals (Tukey HSD test at p-value 0.05). The coefficient of variation (CV) increased with $L$ and decreased with number of cells per simulation (Fig. S5). In all three parameter sets, the CV for *L* and $t_{ss}$ was below 1 and 5% respectively for the number of cells used for all simulations presented in the main text ($N=10^5$).

Next, to assess dependence of the solutions to the time step, we performed a series of simulations with logarithmically increasing values of the time step ($dt = 0.4/2^{(0:6)}$ s) for each of the three sets of chemotaxis parameters used in the tests for *N*. For the time step used in our presented simulations ($dt = 0.1$ s), the deviation from the minimum step tested ($dt = 6.25\cdot10^{-3}$s) was below 10% (Fig. S6).

#### Chemotaxis signalling pathway model

The chemotactic behaviour of the cells in the IBM was simulated with a coarse-grained chemotactic pathway model described extensively elsewhere (Tu et al. 2008; Kalinin et al. 2009; Shimizu et al. 2010). A list and description of all parameters used in the chemotactic pathway models is given in Table S2. Briefly, the chemotaxis signal transduction pathway in *E. coli* has three major components: the transmembrane methyl-accepting chemotaxis protein (MCP) receptors that bind to molecules of the environmental chemoeffector, the flagellar motors, and a pool of different cytosolic proteins that transfer the signal from the MCP to the motors. Essentially, the model predicts the probability of tumbling in response to changes in the environmental concentration of the chemoattractant. It has three dynamic variables: the perceived chemoattractant concentration *C(t)*, the average kinase activity of the MCPs *a(m)* and the average methylation level of the MCPs *m*(*t*) that acts as the memory of the system.

The kinase activity *a(t)* is derived from the Monod-Wyman-Changeux (MWC) model for receptor cooperativity (Mello and Tu 2005; Tu 2013) as:

$$a(t)=\left[1+e^{\left(N_d\left(g_m(m(t))+\ln\left(\frac{1+c(t)/K_i}{1+c(t)/K_a}\right)\right)\right)}\right]^{-1} \qquad \textit{Equation S1}$$

where $N_d = 6$ is the number of dimers of the aspartate sensing receptor (*Tar*) in an all-or-none MWC complex in *E. coli* (Tu 2013), the function $g_m(m(t))$ is the methylation-level dependent free energy difference and $K_I = 18$ µM and $K_A = 2903$ µM are the dissociation constants of chemoattractant molecules to the MWC complex in its inactive and the active states, respectively. The free energy difference is modelled as $g_m(m(t))=\beta(m_0-m(t))$, where $\beta\approx1.7$ (in units of *kT*) is the free-energy change per added methyl group and $m_0\approx1$ (Kalinin et al. 2009).

Following Kalinin et al (Kalinin et al. 2009), we modelled the kinetics of the methylation level with the linear ODE $dm/dt=K_R(1-a(t))-K_B a(t)$, where $K_R=K_B=0.005$ s$^{-1}$ are the rates of methylation for the inactive receptors and of de-methylation for the active receptors, respectively. Although Michaelis-Menten kinetics have been reported for methylation (Shimizu et al. 2010), some preliminary tests showed that results were almost identical, so we used the simpler linear model. At the beginning of a simulation, the methylation state of each cell was set at the adapted state corresponding to the average nutrient concentration at its initial location along the gradient.

##### *Run-and-tumble switch*

The last step in the signalling transduction network of cytosolic proteins is the phosphorylation of the protein CheY. The binding of $CheY_P$ (where the subscript P indicates the phosphorylated state) to a flagellar motor causes the motor to switch from counter-clockwise (CCW) to clockwise (CW) rotation, causing a tumble. The clockwise bias *CWB*, that is the proportion of time in which some of the flagellar motors in a cell are rotating CW and therefore causing a tumble, depends on the cytosolic concentration of $CheY_P$. This dependence follows a Hill equation with a coefficient $H$ = 10.3±1.1 (Cluzel et al. 2000):

$$CWB=\frac{1}{\left(\frac{K_m}{[CheY_p]}\right)^H} \qquad \textit{Equation S2}$$

where $K_m$ is the $CheY_P$ concentration producing a $CWB$ = 0.5.

We assume that the kinase activity of the receptor is proportional to the cytosolic concentration of $CheY_P$, i.e., $a(t)\sim[CheY_P]$, and define $a_{1/2}$ as the kinase activity that induces $CWB$=0.5, so that $a_{1/2}\propto K_m$ (Kalinin et al. 2009). Thus, $\frac{K_m}{[CheY_p]}=\frac{a_{1/2}}{a(t)}$. We further assume that the tumble bias $TB$ = $\tau_t/(\tau_t+\tau_r)$, which we obtained from analyses of individual behaviour in homogeneous environment, is a good proxy for *CWB*. Substituting concentrations ([$CheY_P$] and $K_m$) by activities ($a$ and $a_{1/2}$) and *CWB* by *TB* in Equation S2 yields:

$$\frac{\tau_t}{\tau_t+\tau_r(t)}=\frac{1}{\left(\frac{a_{1/2}}{a(t)}\right)^H+1} \qquad \textit{Equation S3}$$

We solve equation S3 for $\tau_r$(t), whose inverse is the frequency of the Poisson process describing the probability of starting the tumble. Thus, during a time step $\Delta t$, the probability $p$ of a running cell to start tumbling is $p=\frac{\Delta t}{\tau_r(t)}$. However, to compute $\tau_r$(t) at each time step from equation S3 we need estimates for $\tau_t$ and $a_{1/2}$, both of which we assume to be independent of the external ligand concentration (Berg and Brown 1972) but to change with cell length (Guadayol et al. 2017). We obtained cell-dependent estimates for $\tau_t$, (as well as for $\tau_r$ in the absence of chemical gradients, when it can be assumed to be constant) from experiments of elongated *E. coli* swimming in homogeneous medium (Guadayol et al, 2017). To obtain cell-dependent estimates for $a_{1/2}$, we solved equation S3 for $a_{1/2}$ and then used the estimates for $\tau_r$ and $\tau_t$ in homogeneous conditions, assuming $a$ to be, in the absence of chemical gradients, independent of time and constant with a value of 0.5 (Kalinin et al. 2009).

At the beginning of each tumble, its duration was drawn from an exponential distribution with mean $1/\tau_t$ (Berg and Brown 1972).

#### Theoretical background

A framework of hypotheses regarding optimal cell length for chemotaxis can be formulated by combining the empirical functional responses of the main motility parameters to cell elongation in the absence of chemical gradients previously reported by our group (Guadayol et al. 2017) with theoretical models that predict chemotactic performance based on physical constraints to microbial motility (Dusenbery 1998; Locsei 2007; Schuech et al. 2019) or link individual chemotactic behaviours to population distributions (Lovely and Dahlquist 1975; Rivero et al. 1989; Ahmed and Stocker 2008; Tindall et al. 2008). The main conclusions of this review are developed in the following sections and summarized in table S1.

#### Empirical functional responses

The individual motility parameters that have been experimentally shown to be sensitive to cell length in the absence of chemical gradients are: rotational friction coefficient $f_r$, swimming speed $v$, tumble bias $TB$ and directional persistence $\alpha_p$ (Fig. S1, Guadayol et al. 2017). In our experimental system $v$ decreased with increasing cell length in a manner consistent with a cell-size invariant metabolic power. $TB$ increased with cell length as tumble times $\tau_t$ increased and run times $\tau_r$ decreased. Finally, $\alpha_p$ increased as fr increased with cell length, making it more costly for long cells to change orientation during a tumble, but also diminishing the influence of rotational Brownian motion and allowing for straighter runs. In summary, elongation in *E. coli* induces a change in motility pattern from the classical run-and-tumble to a run-and-stop/reverse pattern.

#### Physical constraints

Dusenbery (Dusenbery 1998) examines the effect of cell elongation on swimming efficiency and chemotactic performance on the basis of Perrin's model for hydrodynamic resistance of rigid ellipsoids of revolution at very low Reynolds numbers (Perrin 1934). For swimming efficiency, Dusenbery predicts an optimal axial ratio (i.e., the ratio between long and short axis of the ellipsoid) of 1.952. This value corresponds to an *E. coli* cell length of 1.5 µm. However, the increase in swimming efficiency is less than 5% better than that of a spherical cell. For chemotactic performance, Dusenbery bases his analysis on theoretical changes in the signal-to-noise ratio (S/N) perceived by a bacteria swimming in a linear gradient. This sets an upper limit to chemotactic ability. This approach predicts an increase in S/N ratio (and hence on chemotactic performance) with swimming speed which is in contrast with predictions from transport models (see below). Dusenbery's analysis indicates that S/N is ultimately controlled by frictional coefficients, both translational, through its effects on swimming speed, and rotational, through its effects on running orientation. Thus, $S/N \propto f_r^{3/2}/f_t^{1/2}$, where $f_r$ is the rotational friction coefficient around the minor axis of the ellipsoid of revolution, and $f_t$ is the translation friction coefficient along the major axis (Dusenbery 1998), which leads to an ever-increasing S/N with elongation.

Dusenbery's argument was based solely on translational drag of the cell body. Schuech et al (2019), as well as an earlier study by Shum et al (2010) on ellipsoidal cells, employed much more realistic models of self-propulsion driven by a rotating flagellum, and confirmed Dusenbery's prediction that slightly elongated morphologies swim more efficiently. Schuech et al also predicted chemotactic S/N to increase without bound with elongation, though not as strongly as Dusenbery's prediction. However, Dusenbery and Schuech et al's sole use of S/N to quantify chemotactic performance neglected many other shape-dependent constraints on chemotaxis that we were able to account for here via experimental measurements.

#### Transport models

Chemotaxis is classically modelled using transport models of the Keller-Segel type that characterize chemotactic responses with population-level parameters such as random motility coefficient $\mu$ and chemotactic speed $v_c$ (Keller and Segel 1971). These population parameters can be related to individual-level motility parameters, such a such as swimming speed $v$ and mean length of run (Rivero et al. 1989; Ahmed and Stocker 2008). Thus, given empirical functional relationships of these individual parameters with cell length, it is possible to infer the effect of elongation on chemotactic performance.

Random motility coefficient $\mu$ parametrizes the diffusive dynamics of swimming bacteria at population scales in the absence of any chemical gradient and is analogous to the molecular diffusivity resulting from Brownian motion. It is, in fact, frequently referred to as the bacterial diffusion coefficient. It can be estimated from individual motility parameters in two dimensions as (Ahmed and Stocker 2008):

$$\mu=\frac{16_{2D}^{2}\,\tau_r}{3\pi^2\left(1-\alpha_p\right)} \qquad \text{Equation S4}$$

where $v_{2D}$ is the average 2D swimming speed of bacteria during runs and $\tau_r$is the mean duration of runs in seconds. We may expect either positive or negative responses to elongation in *E. coli*, depending on the relative sensitivity of the individual parameters to cell length.

The chemotactic velocity $v_c$ is the mean speed at which a cell moves up a chemoattractant gradient. In two dimensions it can be calculated as (Ahmed and Stocker 2008):

$$v_c=\frac{8\,v_{2D}}{3\pi}\frac{\left(\tau_r^{+}-\tau_r^{-}\right)}{\left(\tau_r^{+}+\tau_r^{-}\right)} \qquad \text{Equation S5}$$

where $\tau_r^{+}$ and $\tau_r^{-}$ are the mean run times in seconds when bacteria travel up and down the chemical gradient respectively. Setting aside $\tau_r^{+}$ and $\tau_r^{-}$, for which the effect of elongation is unknown, $v_c$ should decrease as *E. coli* cells elongate because of the response of $v_{2D}$. Locsei (Locsei 2007) uses an analytical model to explore the effects of Brownian motion and of $\alpha_p$ on chemotactic velocity $v_c$. In his model, the chemotactic response is modelled phenomenologically with a response function. His results show that $v_c$ peaks at high $\alpha_p$, but rapidly tends to 0 as $\alpha_p$ approaches 1 (that is, when reorientations are virtually inexistent). Experimental data (Fig. S1d) shows that, although $\alpha_p$ increases with cell length, it does not achieve the values for which a collapse in $v_c$ is predicted (Locsei 2007).

The ratio $\mu/v_c$ yields the precision length $L$, which is the parameter we have used to evaluate chemotactic performance:

$$L=\frac{4\,v_{2D}}{\pi\left(1-\alpha_p\right)}\frac{\left(\tau_r^{+}\,\tau_r^{-}\right)}{\left(\tau_r^{+}-\tau_r^{-}\right)} \qquad \text{Equation S6}$$

Both $v_c$ and $\mu$ increase with $v_{2D}$. The net effect at steady state is a linear increase in $L$ with $v_{2D}$. Given the experimentally observed inverse relationship between swimming speed and cell length (Fig. S1b), the expected outcome is therefore a decrease in precision length $L$ as cells elongate. In other words, cell elongation would be expected to help *E. coli* aggregate towards nutrient sources.

Similarly, $\alpha_p$ can affect $L$ in competing ways. The K-S modelling framework predicts an increase in $L$ as cells elongate, whereas Locsei's model predicts the opposite since it predicts $v_c$ increases with $\alpha_p$.

In summary, based on the empirical responses measured by the authors and previous models of chemotaxis, elongation may show contrasting effects on chemotactic performance. Random motility may increase because directional persistence is enhanced, although this effect could be counterbalanced by decreases in speed. Similarly, chemotactic velocity may decrease with cell length due to decreased swimming speed, or alternatively, may increase because of higher directional persistence. Unfortunately, this theoretical framework is incomplete because we do not yet have experimental data on the effect of elongation on the asymmetry between upgradient and downgradient run lengths, which in the case of *E. coli* is what ultimately drives the chemotactic behaviour.

### Supplementary tables

**Table S1**: Summary of the hypothetical responses of motility parameters to elongation based on previously published mechanisms.

| **Parameter** | **Mechanism** | **Favours** | **References** |
|---|---|---|---|
| Translational friction | Slightly elongated cells (aspect ratio ~1.95) minimize translational drag and therefore optimize swimming efficiency. | Intermediate cells | (Dusenbery 1998) |
| Translational friction along and rotational friction about the long axis | High translational friction increases energy expenditure, whereas high rotational friction minimizes energy lost to body counter-rotation. | Intermediate cells | (Schuech et al. 2019) |
| Rotational friction around the short axes | Increasing rotational friction allows for longer runs, as running cells take longer to lose orientation. Longer runs increase differences between chemoattractant measurements, and therefore improve chemotactic signal-to-noise ratio. | Long cells | (Dusenbery 1998; Schuech et al. 2019) |
| Swimming speed | Chemotactic velocity increases linearly with swimming speed while random motility coefficient increases quadratically with speed. The net effect is an increase in precision length. | Long cells | (Rivero et al. 1989) |
| Directional persistence | Chemotactic velocity increases with directional persistence | Long cells | (Locsei 2007) |
| | Restriction of reorientation angles leads to higher random motility coefficients. | Short cells | (Lovely and Dahlquist 1975) |
| Response times (tumble time; run time) | The diffusion time of intracellular signals increases quadratically with cell length. Slower responses lead to longer precision lengths | Short cells | (Segall et al. 1985) |
| | In very long cells, flagella may be too far apart for effective bundling or may form competing bundles of flagella. | Short cells | (Lee et al. 2021) |
| Run time | Flagella become desynchronized in long cells, leading to shorter run lengths, and therefore to lower signal-to-noise ratios | Short cells | (Dusenbery 1998; Maki et al. 2000) |

**Table S2**: List of symbols used in the chemotactic pathway model

| **Symbol** | **Parameter** | **Definition** | **Units** |
|---|---|---|---|
| *C* | Chemoattractant concentration | Environmental concentration of the chemoattractant | µM |
| *CWB* | Clockwise bias | Fraction of time an individual motor spends rotating clockwise | - |
| *a(t)* | Kinase activity | Average kinase activity of the methyl-accepting chemotaxis proteins receptors | - |
| $a_{1/2}$ | Kinase activity at steady state | Average kinase activity that induces a CWB =0.5 | - |
| *m(t)* | Methylation level | Average methylation level of the receptors of the methyl-accepting chemotaxis proteins receptors | - |
| $m_0 = 1$ | Steady state methylation | Average methylation level such that fm(m0) = 0 | - |
| $\beta$~1.7 | Free-energy change | Free-energy change per added methyl group | kT |
| $g_m(m)$ | Free energy difference | $g_m(m(t)) = \beta(m_0 - m(t))$ | kT |
| $N_d$ | Number of dimers | Number of dimers of the aspartate sensing receptor (Tar) in an all-or-none MWC complex in *E. coli* | - |
| $K_i = 18$ | Dissociation constant to inactive receptors | Dissociation constant of aspartate to the inactive transmembrane receptors | µM |
| $K_a = 2903$ | Dissociation constant to active receptors | Dissociation constant of the aspartate to the active transmembrane receptors | µM |
| $K_R = 0.005$ | Methylation rate | Rate of methylation for the inactive transmembrane receptors | $s^{-1}$ |
| $K_B = 0.005$ | De-methylation rate | Rate of de-methylation for the active transmembrane receptors | $s^{-1}$ |
| $K_m$ | Dissociation constant to flagellar motor | Dissociation constant of [CheY-P] to the flagellar motor | µM |
| $H$=10.3±0.1 | Hill coefficient | Coefficient of the Hill equation fit to CWB vs [CheYP] | - |

### Supplementary figures

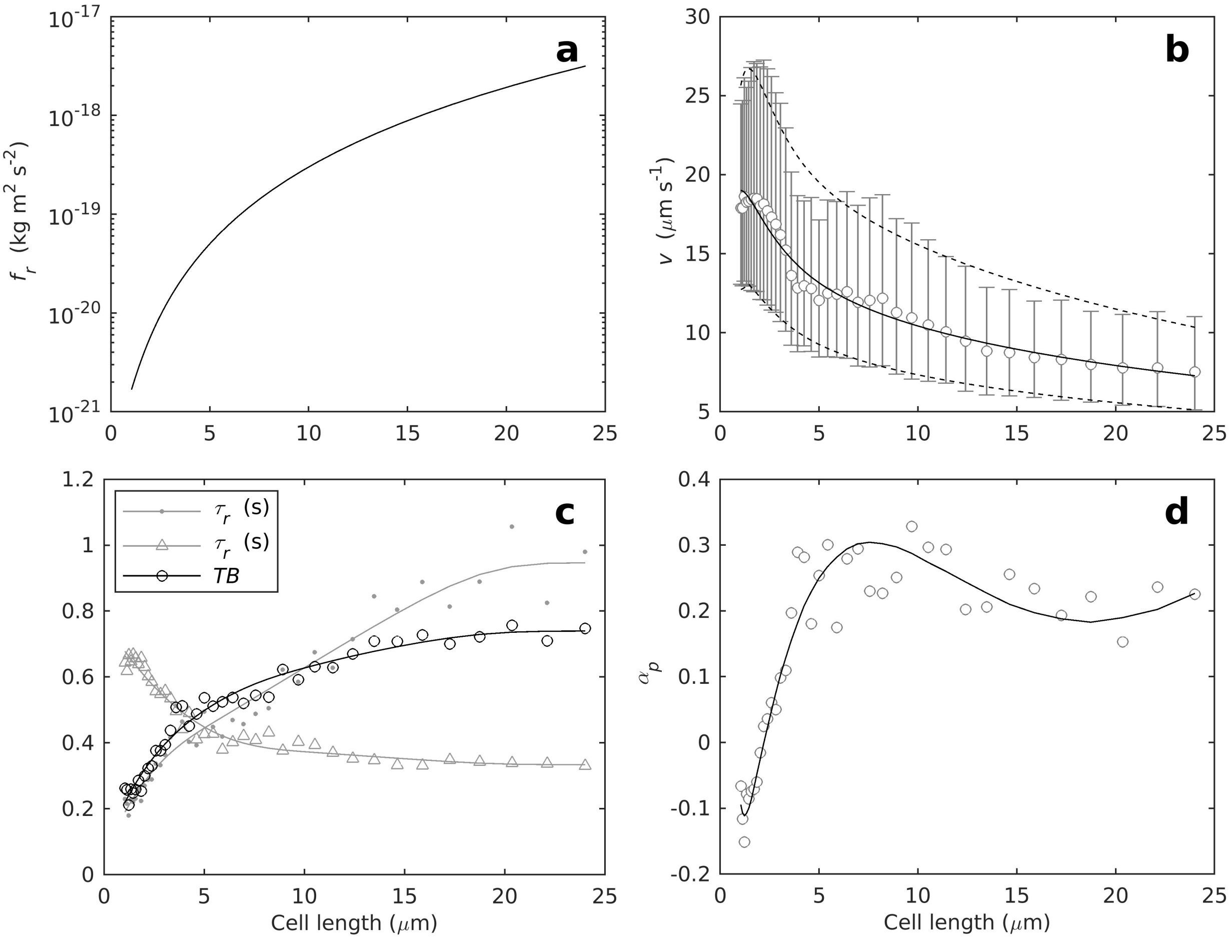


***Figure S1**: Empirical functional responses of the individual motility parameters to cell length used in the IBM simulations. The experimental data, represented by markers, were obtained from Guadayol et al. (2017) dataset of cephalexin-treated* E. coli *cells swimming in a homogeneous environment. Lines are the smoothing functions used in the IBM simulations to model the responses of the motility parameters to cell length (see section "Motility parameters' responses to elongation" in Supplementary methods for details). **a**) Coefficients for rotational friction about the short axes, obtained from the theoretical equations describing rotation of ellipsoids of revolution (Dusenbery 1998). **b**) Average swimming speeds during runs. Circles are empirical averages per cell length class, error bars are the standard deviations, the continuous and dashed lines show the spline function fitted to lognormal distribution parameters per cell-length class used in the IBM simulations. **c**) Average run time ($\tau_r$). tumble time ($\tau_t$) and tumble bias ($TB = \tau_r/(\tau_t + \tau_r)$) per cell class. **d**) Average directional persistence ($\alpha_p$) per cell length.*

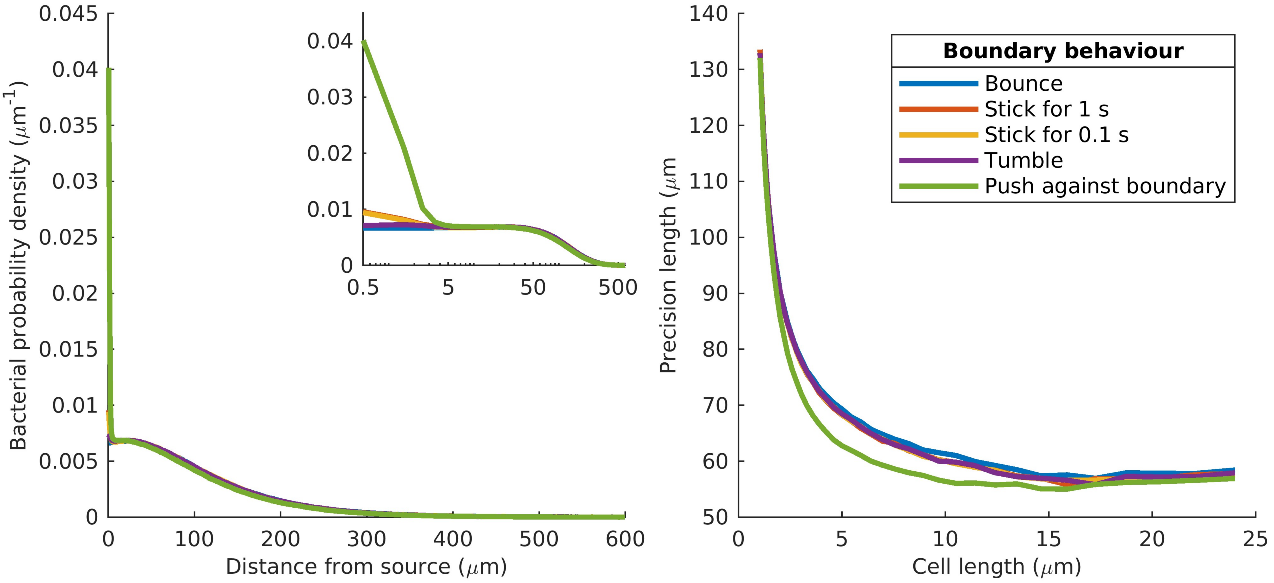


***Figure S2**: Model cell behaviour when encountering walls, described in section “Model behaviour at walls” in Supplementary methods. Left panel shows the steady state distribution of cells across the channel in the presence of a linear gradient of aspartate ($\nabla C/\overline{C} = -1.2\,mm^{-1}$). The inset shows the distributions in a logarithmic x-axis to highlight differences among behaviours close to the walls. Right panel shows steady state L vs cell length for the same gradient of aspartate.*

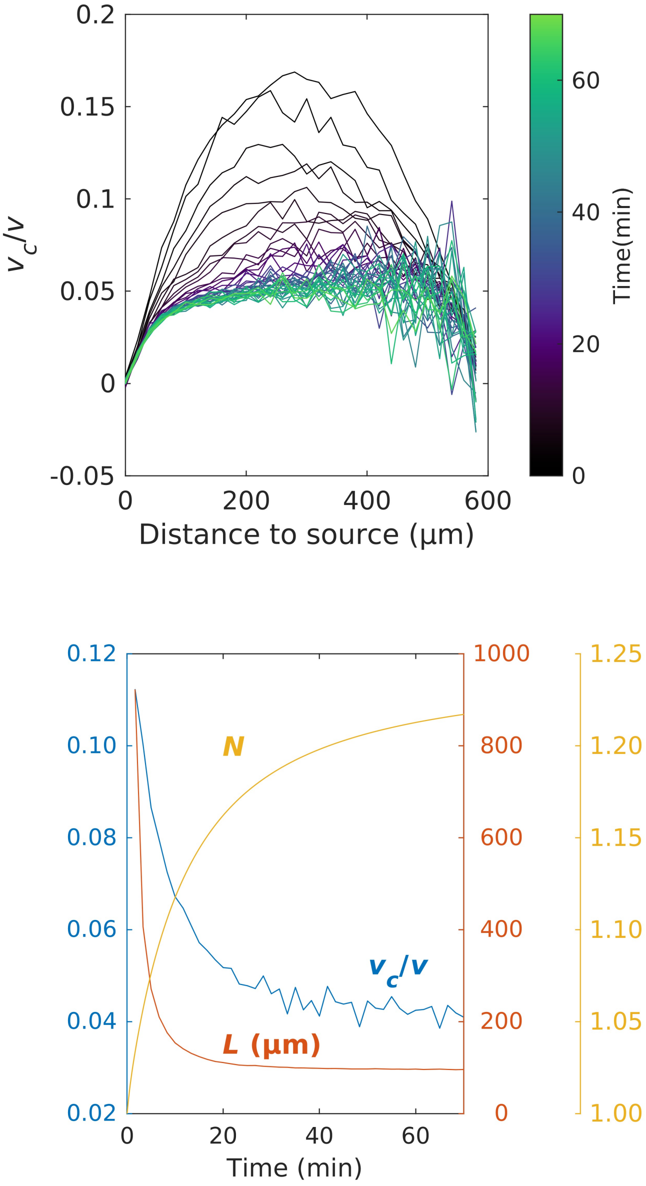


***Figure S3**: Space and time-resolved estimates of chemotactic parameters from simulations of wild-type* E. coli *cells swimming in a linear gradient of aspartate* $(\nabla C/\bar{C} = -1.2\,mm^{-1})$*. Top panel shows normalized chemotactic velocity* ($v_c/v$) *vs distance to nutrient source, colour-coded by time. Bottom panel shows the temporal dynamics of the average* $v_c/v$ *in the centre of the channel (i.e. at distances between 250 and 350 μm), the precision length* L *and the integrated nutrient exposure* N.

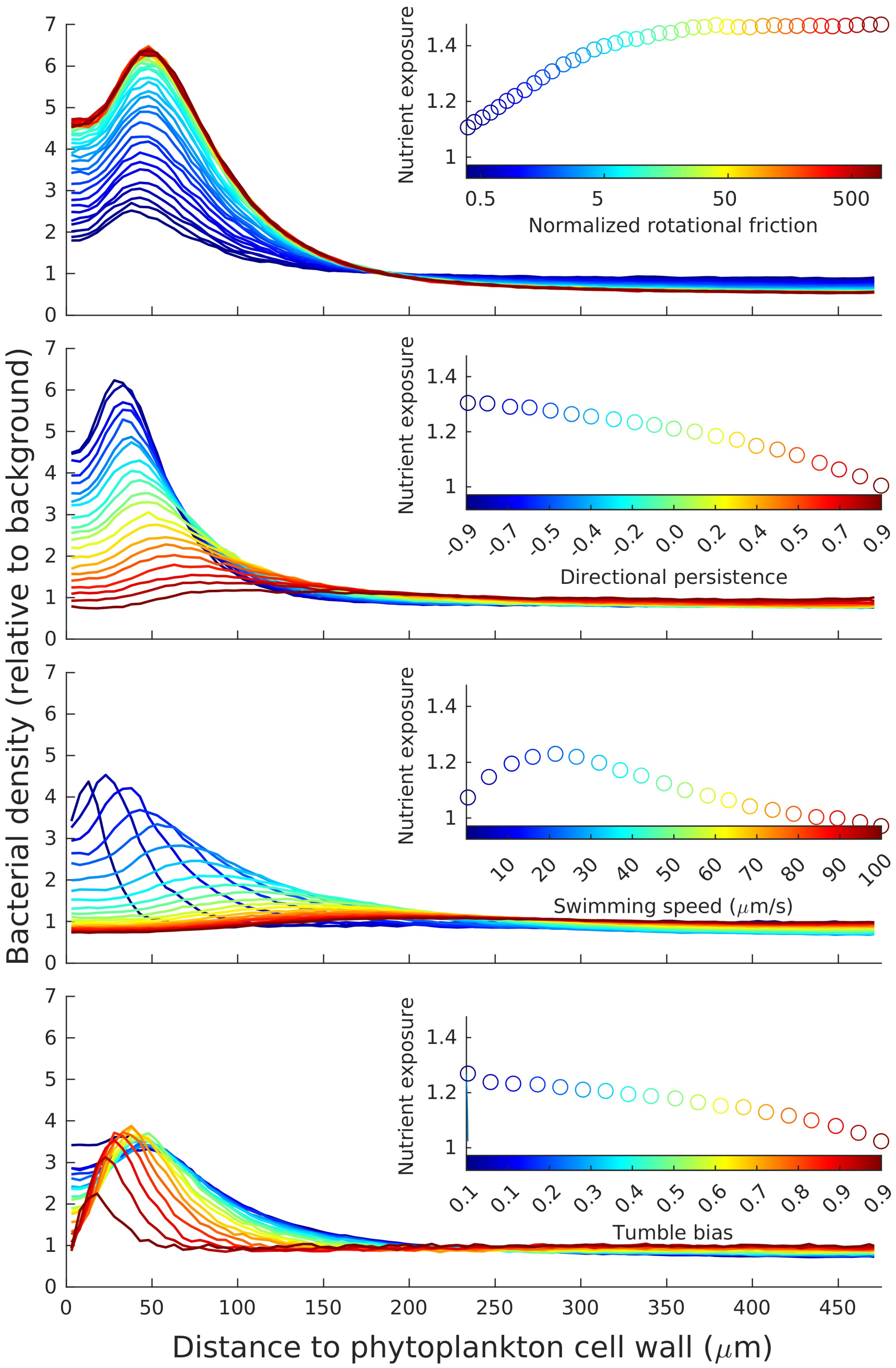


***Figure S4***: *Sensitivity analyses of the phycosphere IBM model for rotational friction (normalized to that of a 1.7 µm long wild-type cell), directional persistence, swimming speed and tumble bias. Main panels show the distribution of bacteria as a function of the distance to the phytoplankton cell length. Subpanels show the colour code for the corresponding main panel and the integrated nutrient exposure normalized by that of a population of uniformly distributed non-chemotactic bacteria.*

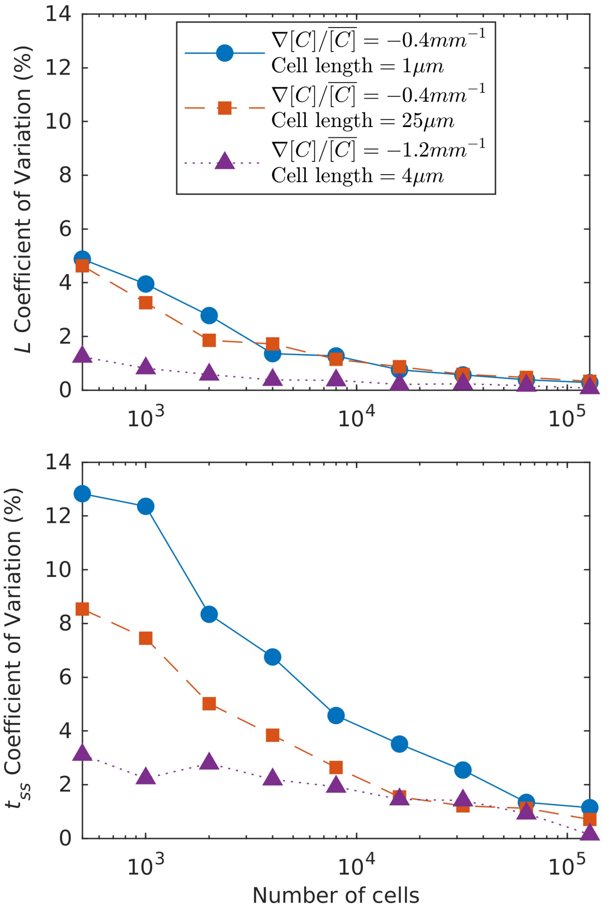


***Figure S5:*** *Convergence test for numbers of cells in the simulation. Three different sets of conditions were used to account for the most extreme values of precision length scale (L) and time to steady state (T) in our experimental setup (see Fig. 2): blue circles show the shallowest gradient* $(\nabla C/\overline{C} = -0.4\,mm^{-1})$ *for the shortest cells (1µm), which resulted in L = 307 µm and a time to steady state* $t_{ss}$ *= 63 min; red squares show simulations with the shallowest gradient* $(\nabla C/\overline{C} = -0.4\,mm^{-1})$ *for the longest cells (25µm), which gave L= 135 µm and* $t_{ss}$ *= 293 min; Purple triangles show simulations with the steepest gradient* $(\nabla C/\overline{C} = -1.2\,mm^{-1})$ *and a cell length of 4 µm, which gave L= 61 µm and* $t_{ss}$ *= 18 min.*

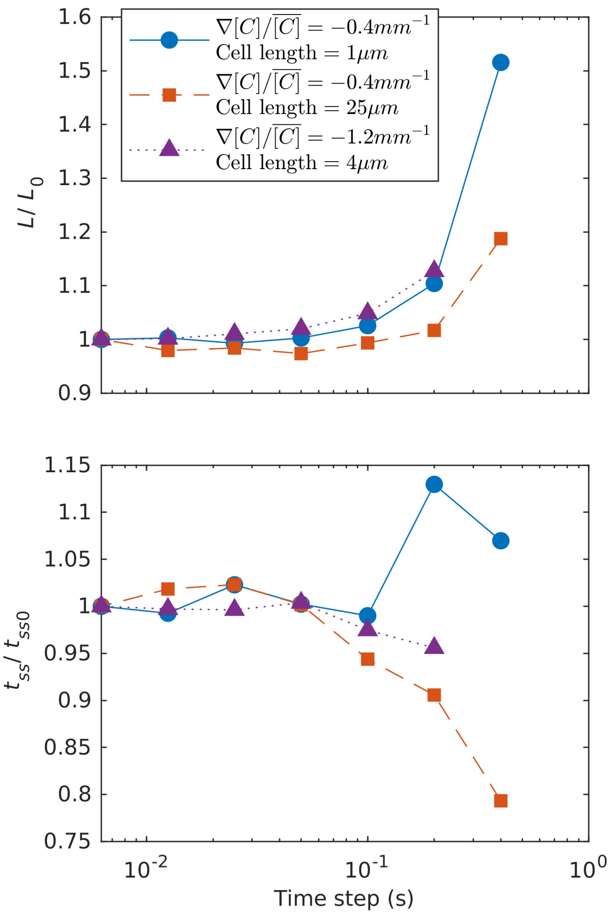


***Figure S6:*** *Time step test for the same three sets of conditions used for the convergence test (Fig. S5). Top plot shows L vs time step; bottom plot shows* $t_{ss}$*. Values are normalized by shortest time step values. Symbols as in Fig. S5*
